# From perception to valence: a pair of interneurons that assign positive valence to sweet sensation in *Drosophila*

**DOI:** 10.1101/2025.10.31.685871

**Authors:** Kevin William Christie, Tarandeep Singh Dadyala, Phuong Chung, Masayoshi Ito, Lisha Shao

## Abstract

Assigning valence—appeal or aversion—to gustatory stimuli and relaying it to higher-order brain regions to guide flexible behaviors is crucial to survival. Yet the neural circuit that transforms gustatory input into motivationally relevant signals remains poorly defined in any model system. In Drosophila melanogaster, substantial progress has been made in mapping the sensorimotor pathway for feeding and the architecture of the dopaminergic reinforcement system. However, where and how valence is first assigned to a taste has long been a mystery. Here, we identified a pair of subesophageal zone interneurons in Drosophila, termed fox, that impart positive valence to sweet taste and convey this signal to the mushroom body, the fly’s associative learning center. We show that fox neuron activity is necessary and sufficient to drive appetitive behaviors and can override a tastant’s intrinsic valence without impairing taste quality discrimination. Furthermore, fox neurons transmit the positive valence to specific dopaminergic neurons that mediate appetitive memory formation. Our findings reveal a circuit mechanism that transforms sweet sensation into a reinforcing signal to support learned sugar responses. The fox neurons exhibit a convergent–divergent “hourglass” circuit motif, acting as a bottleneck for valence assignment and distributing motivational signals to higher-order centers. This architecture confers both robustness and flexibility in reward processing—an organizational principle that may generalize across species.

## Introduction

How sensory information is transformed into emotionally or motivationally charged representations is a central question in neuroscience: Assigning proper valence––appeal or averseness––to sensory stimuli is fundamental for an organism’s survival, guiding behaviors such as approach or avoidance, learning, and decision-making. Yet the neural circuits that orchestrate valence assignment across brain regions are unknown or poorly understood.

The circuitry that processes taste exemplifies this complexity, because perceived quality and assigned valence often diverge. Sweet taste, for instance, does not invariably signal a rewarding outcome. Thus, determining whether a taste is "good", "neutral", or "bad" likely requires circuits that integrate context, internal state, and experience—rather than a fixed labeled-line system. Consistent with this view, emerging evidence indicates that gustatory processing occurs through multiple layers––the early layers encode taste quality (e.g., sweet), whereas downstream layers assign and update valence after integrating the animal’s internal state and prior experience^1–4^. These architectures likely contain divergent and convergent motifs, with specific nodes acting as hubs to route sensory signals into distinct behavioral responses– ranging from reflexive feeding to reinforcement learning^5,6^.

In *Drosophila*, flexible modification of taste-driven behaviors provides compelling examples of these principles. Fruit flies avoid sugars previously paired with an aversive stimulus, demonstrating learned aversion^7^. Conversely, flies bearing gut tumors––a drastic alteration in internal state– –show reduced aversion to a bitter compound with antitumorigenic properties^8^. These observations indicate that tastant valence can be dynamically altered by mechanisms acting downstream of the primary gustatory neurons. Nevertheless, it remains unclear where along the gustatory pathway the sensory quality, i.e., “sweet”, is first converted into a positive valence, i.e., “good”.

To elucidate where and how an innate valence is assigned and modified to produce flexible behavioral outputs, we must identify the intermediate neuronal layers responsible for transforming sensory quality into a hedonic signal. Operationally, neurons assigning the valence of sensory stimuli are typically necessary and sufficient for sensory-driven behavioral responses––innate attraction/aversion or learned preference/aversion––without impairing sensory identity recognition, and capable of overriding the intrinsic valence of a stimulus^1,5,6,9^. These criteria guide the search for interneurons that implement valence assignment.

Here, we combine connectome mapping with targeted functional interrogation in the fruit fly, *Drosophila melanogaster,* to pinpoint where the positive valence of tastants first arises within the taste circuitry and how it is relayed to downstream learning circuits. We identified a pair of subesophageal zone (SEZ) neurons that assign a positive valence to the activity of sweet gustatory neurons. Activating these SEZ neurons is sufficient to drive appetitive behaviors, including approach, proboscis extension, and increased consumption of both food and water.

Crucially, these neurons differ from the sweet gustatory receptor neurons and downstream sensorimotor interneurons in two respects: First, their activation, when paired with a neutral odor cue, drives appetitive olfactory learning. Second, their activation can override a tastant’s intrinsic valence, rendering neutral or even aversive tastes attractive. Together, we propose a mechanism by which positive valence is attached to sweet taste inputs and subsequently routed into associative learning networks.

## Results

We started searching for neurons in flies whose activation has a valence––rewarding or punishing––by screening 241 restricted-expression split-GAL4 drivers ^10^ (Figures 1A and S1A) using an optogenetic positional preference assay in which flies choose to prefer or avoid optogenetic neural activation^11^. One of the positive hits labeled the PPM3 cluster of dopaminergic neurons (DANs) and a pair of neurons that innervate the SEZ (Figure S2A). This piqued our interest because dopaminergic neurons that innervate the mushroom body have been implicated as a key hub for valence processing, analogous to the mammalian cerebellum^12,13^. Unexpectedly, however, further characterization attributed the positive positional preference to the SEZ neurons, which were previously named fox and have unknown function^14^ (Figures 1B-D and S3). Along with two more specific drivers (Figure 1C), we found that optogenetically activating fox neurons induced positional preference, while optogenetic suppression^15^ of fox induced positional aversion (Figures 1B and S2B).

**Figure 1.**
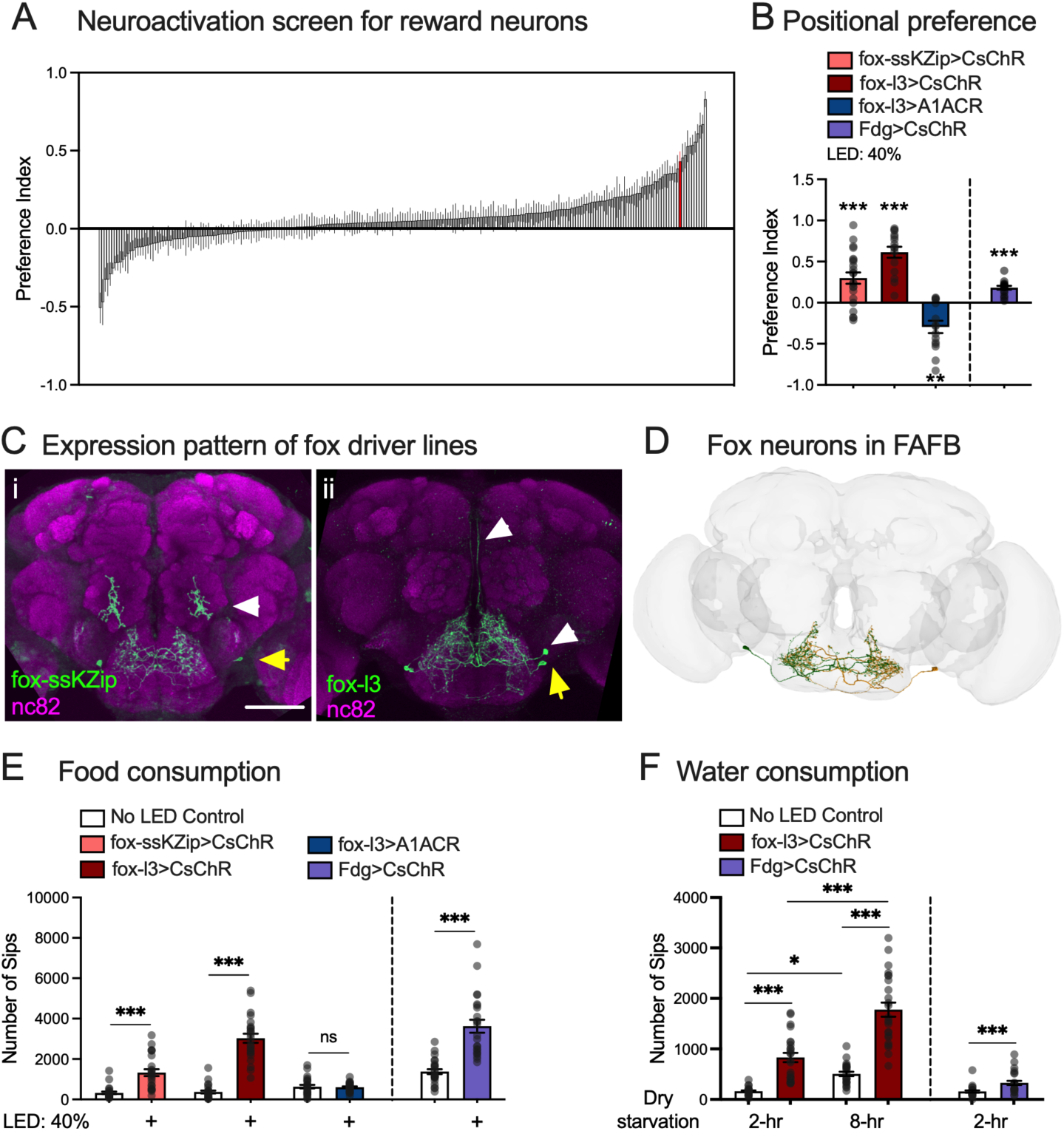
Activating a pair of SEZ interneurons, called fox, is sufficient to drive appetitive behaviors in flies. A. Results of a screen for neurons whose activation is rewarding or aversive to flies using a positional preference assay. A preference index greater than 0 indicates attraction and lower than 0 indicates avoidance. Data from female flies are shown. The performance of fox-ss1>CsChrimson in the screen is highlighted in red. N = 10 for each driver line. Each replicate contains 6-8 flies. B. Flies preferred the illuminated side in the positional preference assay when their fox neurons were optogenetically activated by CsChrimson driven by fox-ssKZip (fox-ss1 + TH-D-KZip; excludes PPM3 neurons by expressing the ’killer zip’ in dopaminergic neurons (TH-D-KZip^32^)), and fox-ss2-13 (fox-l3, a new split-GAL4 driver^14^). Both drivers isolated the same pair of neurons in the sube-sophageal zone, identified as fox neurons. Flies avoided the illuminated side when their fox neurons were blocked by the chloride transporter A1ACR. Like fox neurons, activating a command neuron for feeding, the third-order gustatory interneuron Fdg neurons (Fdg>CsChrimson), induced positional preference. N = 22 for fox-ssKZip>CsChrimson; N = 15 for fox-l3>CsChrimson; N = 15 for fox-l3>A1ACR; N = 15 for Fdg>CsChrimson. Each replicate contains 6-8 flies. One-sample *t* test (hypothetical value = 0), ** p<0.01, *** p<0.001. C. Immunostaining of CsChrimson-mVenus driven by fox-ssKZip and fox-l3 driver lines that label fox neurons: (i) fox-ssKZip); (ii) fox-l3. Yellow arrows indicate fox neurons; white arrows indicate non-fox neurons labeled by the same driver. mVenus signal and neuropils were revealed by antiGFP and nc82 staining, respectively. Scale bar: 50 μm. D. Fox neurons’ morphology in the female adult fly brain. E. Activating fox neurons with CsChrimson using fox-ssKZip and fox-l3 driver lines increased food consumption in fed flies; however, blocking fox neurons with A1ACR did not affect feeding. Similarly, activating Fdg neurons (Fdg>CsChrimson) increased food consumption in fed flies. N = 24 flies for each group. *t* test (no LED and LED groups of the same genotype), *** p<0.001. F. Activating fox neurons (fox-l3>CsChrimson) significantly increased water consumption when flies were dry-starved for 2 hours and 8 hours. N = 24 for each group. One-way ANOVA with Tukey’s multiple comparison test, ns. P>0.05, ** p<0.01, *** p<0.001. Activating the third-order gustatory interneuron Fdg neurons (Fdg>CsChrimson) also increased water consumption in flies with 2-hr dry-starvation. N = 24 flies for each group. *t* test (no LED and LED groups of the same genotype), *** p<0.001.

## Fox activation drives appetitive behaviors

Fox’s SEZ location prompted us to test its function in feeding-related behaviors. We found that activating fox neurons significantly promotes proboscis extension reflex (PER) (Figure S4) and food consumption^16,17^ (Figures 1E, S1C, and S2C). In contrast, blocking fox neurons, either acutely or chronically, did not significantly affect food consumption (Figures 1E and S5). Intriguingly, activating fox neurons also promoted consumption of water, although intake nevertheless remained regulated by the duration of dehydration (Figure 1F). Thus, the activation of fox neurons drives appetitive behaviors, including approaching and preference for fox neuron activation and increased consumption for food and water.

Fox neurons’ phenotypes resemble those of the Feeding (Fdg) neurons, which function as command neurons for feeding behavior^18^ and account for 10% of the total synaptic inputs to fox neurons. Like fox neurons, activation of Fdg neurons increased food and water consumption and induced acute preference for optogenetic activation (purple bars in Figures 1B, E, F). This parallel, together with fox-driven PER (Figure S4), supports the idea that fox neurons also contribute to the regulation of the feeding motor program. Connectome analysis showed that, although fox neurons were not previously classified as higher-order gustatory interneurons (GINs) ^3,19^ or SEZ local neurons (SEZ-LN)^4^, they receive synaptic inputs from key second- and third-order GINs and project onto premotor neurons controlling PER^19–22^(Figure 2A). Moreover, fox neurons are a major upstream partner for the drosulfakinin neurons (DSKMP1B) in SEZ, accounting for over 20% of DSKMP1B synaptic inputs from SEZ. Because fox neurons both drive appetitive behaviors and interface with the core feeding pathway and internal-state regulators^23^, we next investigated their function in valence regulation, particularly in the context of sweet taste.

**Figure 2.**
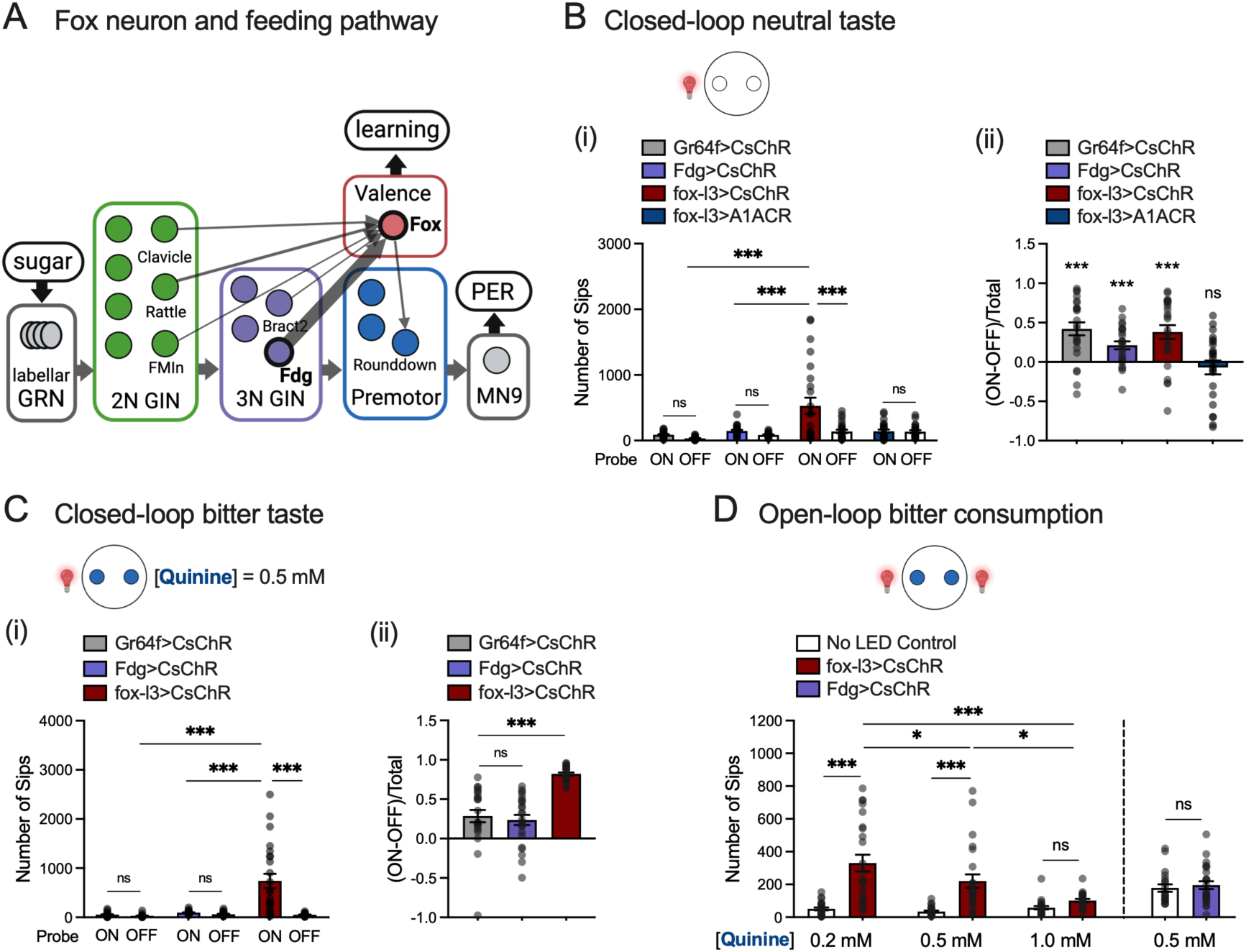
Activation of fox neurons overrides the intrinsic valence of neutral and aversive taste. A. Illustration of fox neural connections with the neural pathways regulating proboscis extension reflex (PER). Synapses from Fdg neurons comprise 10% of the synaptic inputs to fox neurons. B. Optogenetic activation of either Gr64f-expressing GRNs (Gr64f>CsChR), Fdg neurons (Fdg>CsChR), or fox neurons (fox-l3>CsChR) when the fly is feeding on one of the two probes containing agar significantly increased fly’s feeding from the associated probe, whereas optogenetic silencing of fox neurons (fox-l3>A1ACR) with a probe did not change the fly’s choice. Panel (i) shows the number of sips on each probe. One-way ANOVA with Tukey’s multiple comparison test, ns. p>0.05; *** p<0.001. Panel (ii) shows preference, calculated as the difference between sips on associated versus non-associated probe, normalized by the total number of sips ((ON-OFF)/Total). One sample *t* test (hypothetical value = 0), *** p<0.001, ns p>0.05. N = 24 flies for each group. C. Flies’ feeding from one of the two probes containing quinine (0.5 mM) was significantly increased when the probe was associated with fox activation (fox-l3> CsChrimson), but not with Gr64f-expressing GRN (Gr64f> CsChrimson) or with Fdg neuron (Fdg> CsChrimson) activation. Panel (i) shows the number of sips on each probe. One-way ANOVA with Tukey’s multiple comparison test, ns. p>0.05; *** p<0.001. Panel (ii) shows preference, calculated as the difference between sips on associated versus non-associated probe, normalized by the total number of sips ((ON-OFF)/Total). One-way ANOVA with Tukey’s multiple comparison test, *** p<0.001, ns. p>0.05. N = 24 flies for each group. D. Activating fox neurons (fox-l3>CsChrimson) significantly increased flies’ consumption of agar containing 0.2 mM and 0.5 mM quinine but did not promote consumption for agar containing 1.0 mM quinine. N=24 for each group. One-way ANOVA with Tukey’s multiple comparison test, ns. p>0.05, * p<0.05, ** p<0.01, *** p<0.001. In contrast, activating Fdg neurons (Fdg>CsChrimson) did not increase the consumption of quinine (0.5 mM). N = 24 flies for each group. *t* test, ns. p>0.05.

## Fox activation imposes a positive valence on taste

To determine whether fox neurons are sufficient to impose a positive valence on taste, we performed a closed-loop optogenetic feeding assay^17^, where flies chose between two probes containing a neutral tastant (agar), with optogenetic neuronal activation contingent on feeding from one of them. We found that, compared to the unpaired probe, flies exhibited significantly increased preference for the probe associated with activation of Gr64f, Fdg, or fox neurons (Figure 2B), also quantified as the difference between sips on associated versus non-associated probe, normalized by the total number of sips ((ON-OFF)/Total) (Figure 2B, ii). Associating the probe with optogenetic silencing of fox neurons, however, did not cause a preference for either probe (blue bars in Figure 2B). These results demonstrate that activation of fox neurons is sufficient to impose a positive valence on neutral tastants.

Next, we tested whether fox activation could override aversive valence. When we replaced the neutral tastant with a bitter substance (0.5 mM quinine in agar), optogenetic activation of fox still drove a significant increase in consumption from the paired probe (Figure 2C). In contrast, activating Gr64f or Fdg neurons yielded only marginal preference (Figure 2C). Finally, we asked if activating fox neurons promoted overall bitter consumption. We found that the effect of fox-induced consumption scaled inversely with quinine concentration (Figure 2D). This graded response suggests that fox neurons likely function downstream of the sensorimotor feeding pathway to modulate hedonic value rather than simply driving a compulsive feeding behavior (as, e.g., the Fdg neurons). Together, fox activation overrides the inherent neutral or negative valence of tastants, indicating fox neurons play a role in assigning positive valence to gustatory stimuli.

## Fox silencing impairs taste valence without altering taste identity

To assess taste-driven behaviors in the absence of fox function, we silenced fox neurons and measured feeding responses. Fox silencing abolished the increase in food consumption normally elicited by activating Gr64f-expressing neurons (Figure 3A). When we examined preference for the probe paired with Gr64f activation in the closed-loop assay, blocking fox did not change overall choice (Figure 3B, ii), but it significantly reduced the number of sips from the activation-paired probe to levels indistinguishable from the unpaired probe (Figure 3B, i). These findings indicate that silencing fox neurons impairs the appetitive responses driven by sweet-GRN activation.

**Figure 3.**
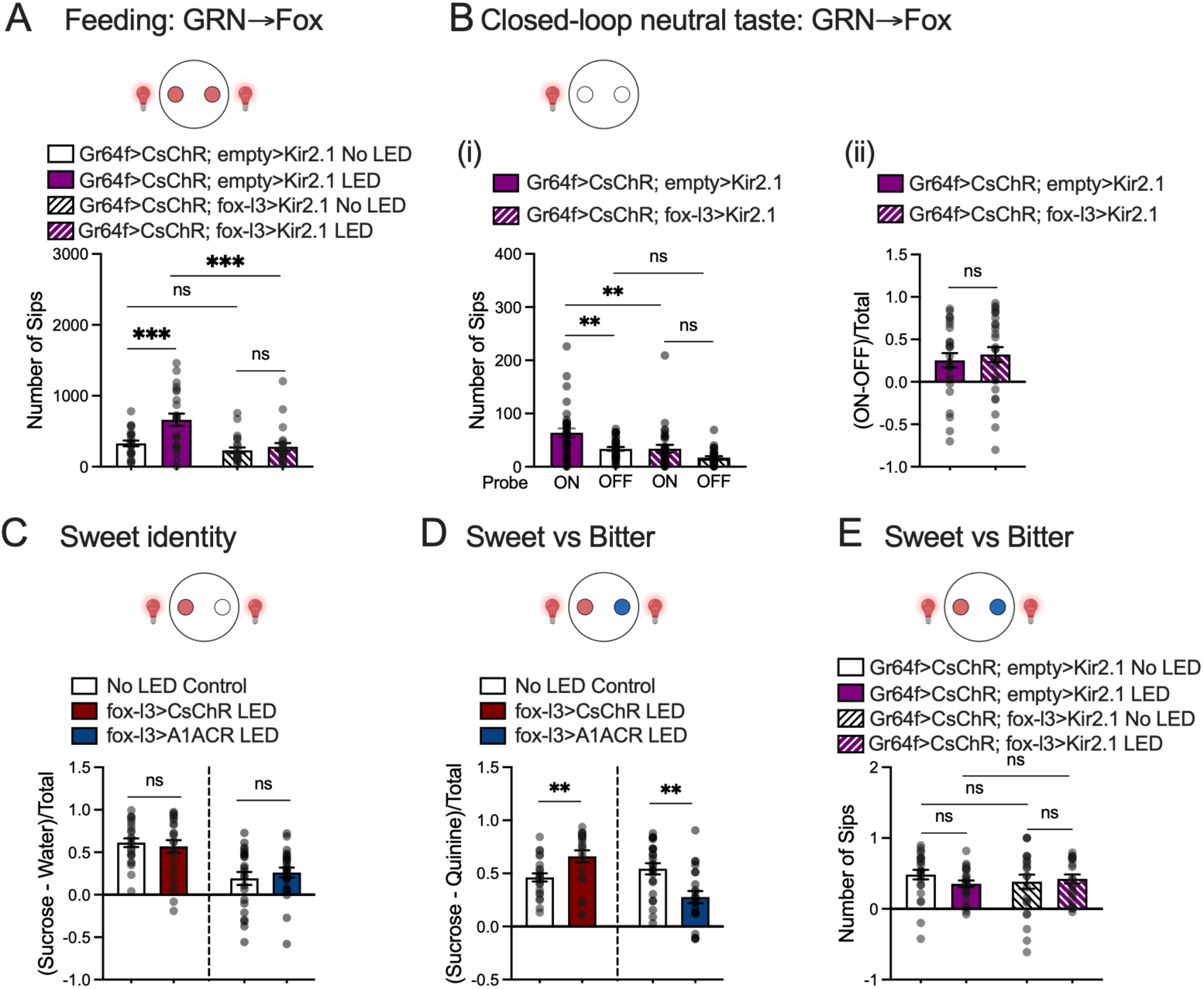
Silencing Fox neurons impairs taste valence without changing recognition of taste identity. A. Optogenetic activation of Gr64f neurons (Gr64f-LexA>LexAop-CsChrimson; empty>UAS-Kir2.1) promoted feeding. In contrast, blocking fox neurons with Kir2.1 (Gr64f-LexA>LexAop-CsChrimson; fox-l3>UAS-Kir2.1) suppressed the enhanced feeding caused by Gr64f activation. N = 24 flies for each group. One-way ANOVA with Tukey’s multiple comparison test, ns. p>0.05; *** p<0.001. B. Activating Gr64f neurons (Gr64f-LexA>LexAop-CsChrimson; empty>UAS-Kir2.1) increased feeding from the agar probe paired with activation, whereas silencing fox neurons (Gr64f-LexA>LexAop-CsChrimson; fox-l3>UAS-Kir2.1) abolished this increase (i). One-way ANOVA with Tukey’s multiple comparison test, ns. p>0.05; ** p<0.01. Flies showed a similar preference for the associated probe with or without fox silencing, preference calculated (ON-OFF)/Total (ii). *t* test, ns (p>0.05). N = 24 flies for each group. C. Optogenetically activating (fox-l3>CsChrimson) or silencing (fox-l3>A1ACR) fox neurons did not affect flies’ ability to distinguish sucrose from agar. [sucrose] = 100 mM. N = 24 flies for fox activation groups and N = 23 flies for fox silencing groups. *t* test, ns (p>0.05). D. Optogenetically activating fox neurons (fox-l3>CsChrimson) increased preference for sucrose over quinine, while blocking fox neurons (fox-l3>A1ACR) decreased the preference. [sucrose] = 100 mM, [quinine] = 0.5 mM. N = 23 flies for each group. *t* test, ns (p>0.05). E. Optogenetic activation of Gr64f-expressing sweet gustatory receptor neurons with (Gr64f-LexA>LexAop-CsChrimson; fox-l3>UAS-Kir2.1, N = 24 flies) or without silencing fox neurons (Gr64f-LexA>LexAop-CsChrimson; empty>UAS-Kir2.1, N=23 flies for no LED control and N=22 flies for LED group) did not change flies’ preference for sucrose over quinine. One-way ANOVA with Tukey’s multiple comparison test, ns (p>0.05).

Such impairment might arise either from a failure to assign valence or from a deficit in taste identity. To distinguish these possibilities, we reasoned that if taste identity and valence are encoded in separate circuits—GRN-GIN for identity and fox neurons for valence—then flies lacking fox activity should still discriminate tastant identity despite lacking proper hedonic valuation. Indeed, fox-silenced flies continued to differentiate sweet from neutral tastants (Figures 3C). Specifically, when choosing between sucrose and agar, neither activation nor silencing of fox neurons altered flies’ preference for sucrose compared to no-LED controls (Figure 3C). As expected, when choosing between sucrose and quinine, fox activation enhanced sucrose preference, whereas fox silencing reduced it (Figure 3D). By contrast, activation of sweet-GRNs did not change fly’s preference for sucrose over quinine with or without silencing fox neurons (Figure 3E). Crucially, fox-silenced flies still retained a net preference for sucrose and aversion to quinine, confirming intact detection of sweetness and bitterness despite disrupted hedonic processing (Figure 3D). Together, these results demonstrate that fox neurons are necessary and sufficient to drive positive, valence-specific feeding behaviors, whereas the GRN-GIN pathway independently encodes tastant identity.

## Fox neurons are essential for appetitive learning induced by GRN activation

In a standard associative learning paradigm, flies learn to associate a neutral odor (conditioned stimulus, CS) with a rewarding sweet tastant (unconditioned stimulus, US)^24^, and the strength of the association can be measured quantitatively by computing the proportion of flies staying in proximity to the source of the paired odor. In a variation of this assay, optogenetic activation of sweet GRNs can substitute for natural sweet tastants, serving as an artificial US to drive learning^25^. This substitution approach allows us to probe how specific neurons contribute to associative memory formation. We hypothesized that fox neurons act as a critical node in generating positive valence and subsequently conveying it to the brain’s associative learning center. Consistent with this, pairing an odor with optogenetic activation of fox neurons produced robust appetitive memory (Figure 4A), whereas optogenetic suppression of fox––despite eliciting acuate aversion (Figure 1B)––did not form an aversive memory in the learning assay (Figure 4A). Notably, activating Fdg neurons failed to generate appetitive learning (purple bar in Figure 4A). Moreover, none of the key GINs that connect to fox––Clavicle, Rattle, FMIn, Bract, Fdg, and Rounddown^19^––generated appetitive olfactory memory when we paired their activation with odors^3^.

**Figure 4.**
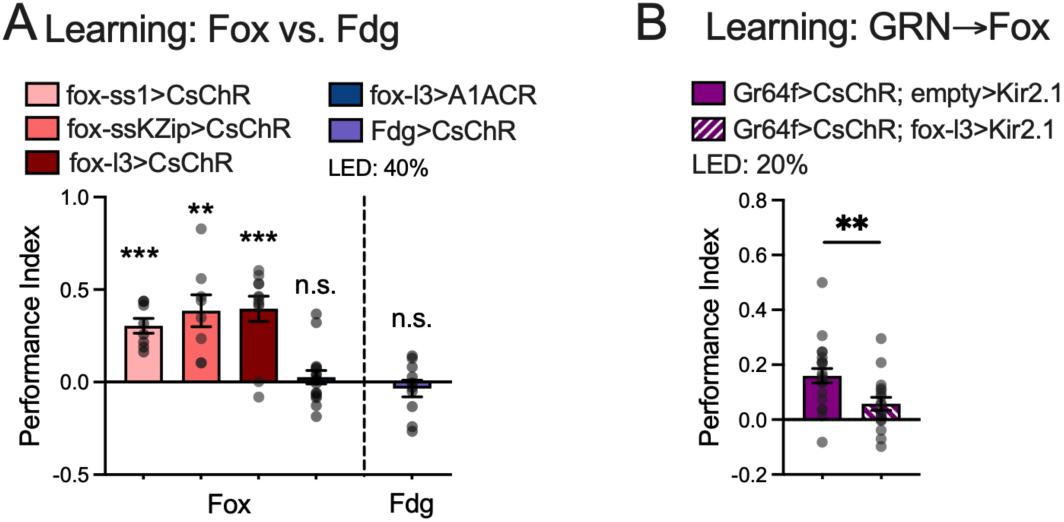
Fox neurons are essential for appetitive learning induced by GRN activation A. Flies formed appetitive memory with fox activation by fox-ss, fox-ssKZip and fox-l3 drivers. Blocking fox neurons resulted in no memory formation. Unlike fox neurons, pairing the activation of Fdg neurons (Fdg>CsChrimson) with odors did not generate appetitive or aversive learning. N = 8 for fox-ss>CsChrimson and fox-ssKZip>CsChrimson; N = 11 for fox-l3>CsChrimson; N = 16 for fox-l3>A1ACR; N = 20 for Fdg>CsChrimson. Each replicate contains 12-14 flies. One-sample *t* test (hypothetical value = 0), ** p<0.01, *** p<0.001, ns. p>0.05. B. Optogenetic activation of Gr64f-expressing sweet gustatory receptor neurons (Gr64f-LexA>LexAop-CsChrimson; empty>UAS-Kir2.1) formed appetitive memory when paired with odors. In contrast, blocking fox neurons with Kir2.1 (Gr64f-LexA>LexAop-CsChrimson; fox-l3>UAS-Kir2.1) suppressed the learning (h, N = 21 and 17) formed by Gr64f activation. Each replicate contains 12-14 flies. *t* test, ** p<0.01.

The results of our learning experiments differentiate fox neurons from the sensorimotor pathway for feeding in the context of associative learning. To further test if fox neurons mediate the valence of GRN activation in appetitive learning, we activated the sweet-sensing GRNs that express Gr64f while simultaneously blocking fox neural activity. We found that blocking fox neurons significantly compromised the learning generated by Gr64f-GRN activation (Figure 4B). Thus, fox neurons are essential for relaying the positive valence of sweet GRN activity to the learning center.

## Fox neurons connect to the learning center via ascending downstream neurons

Our results raise the question of how tastants’ valence is assigned and relayed to the central brain, particularly the learning center. For example, it is well characterized that appetitive gustatory inputs activate the Protocerebral Anterior Medial cluster of dopaminergic neurons (PAM-DANs) that innervate the horizontal lobes of the mushroom body (MB). The changes in PAM-DANs’ activity in turn lead to behavioral changes by modifying the synaptic plasticity between mushroom body intrinsic neurons and output neurons, in a compartment-specific manner^12,13,26,27^. Fox neurons connect to PAM-DANs through 11 ascending neuronal pairs (fox downstream ascending–– FDA––neurons, Figure 5A; full list in Table S1A). Three pairs of FDA neurons synapse directly onto PAM-DANs (FDA Neuron-I) and previously have been shown to be critical for appetitive learning^28^; the other pairs are one synapse upstream of PAM-DANs (FDA neuron-II) and account for 4.4% and 10.5% of total fox synaptic output, respectively.

**Figure 5.**
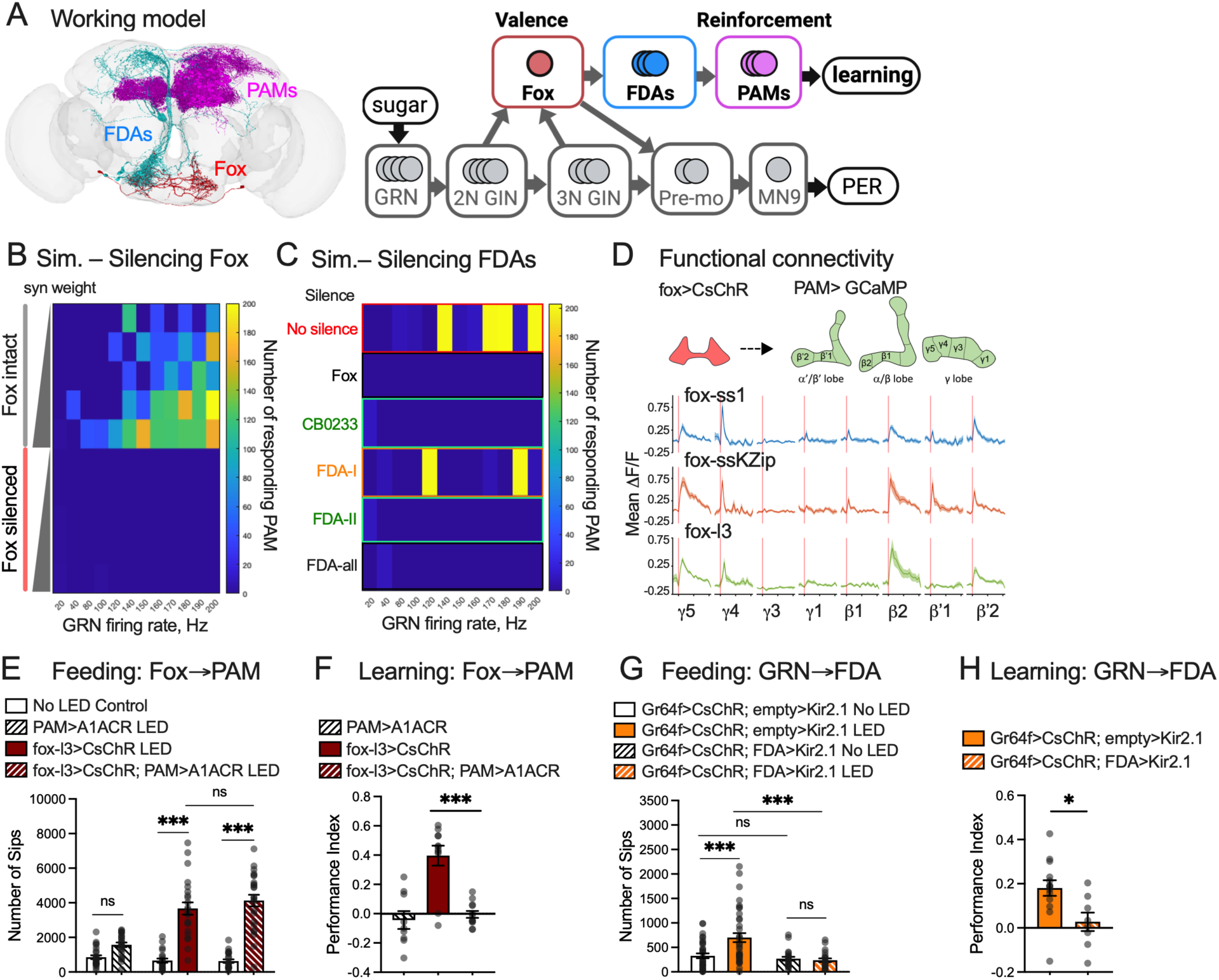
Fox neurons relay GRN signals to PAM dopaminergic neurons. A. Working model. Fox neurons assign valence to GRN-GIN neural activities and convey the valence to PAM through Fox Downstream Ascending (FDA) neurons. B. Heatmap of the number of PAM-DANs that respond to the stimulation of unilateral GRNs and GINs, with and without fox neurons blocked, at various model synaptic weights (0.37, 0.38, 0.382, 0.386, 0.39) and GRN firing rates (20, 40, 80, 100, 120, 140, 150, 160, 170, 180, 190, 200 Hz) in computational simulation. Each number of responsive neurons is an average of five simulation trials with the same parameters. C. Heatmap of the number of PAM-DANs that respond to the stimulation of unilateral GRNs and GINs, with and without fox and FDAs blocked (model synaptic weight = 0.39). FDA-I neurons (orange) synapse directly onto PAM-DANs. FDA-II neurons (green) do not synapse directly on PAM-DANs. CB0233 is an FDA-II cell type with the most synapses with fox neurons. D. Upper panel: Illustration of the functional connectivity experiment. We activated fox neurons while collecting calcium signals from the axons of PAM-DANs that innervate various mushroom body compartments. Lower panel: Optogenetically stimulating fox neurons with three fox drivers led to compartment-specific activation in PAM dopaminergic neurons. Genotype: fox-driver>UAS-Chrimson88-tdTomato; PAM (58E02)-LexA>LexAop-GCaMP7s. N =13 for fox-ss1, N=10 for fox-ssKZip, and N=7 for fox-l3. E-F. Optogenetic activation of fox neurons while blocking PAM dopaminergic neurons with A1ACR (fox-l3>UAS-CsChrimson; 58E02-LexA>LexAop-A1ACR) did not affect feeding (e, N = 24 flies for all groups) but abolished appetitive learning (f, N = 9, 12, 12. Each replicate contains 12-14 flies). One-way ANOVA with Tukey’s multiple comparison test, ns. p>0.05, *** p<0.001. G-H. Blocking CB0233 neurons (FDA-II) with Kir2.1 (Gr64f-LexA>LexAop-CsChrimson; FDA-cb0233>UAS-Kir2.1) suppressed the enhanced feeding (g, N=24 flies for all groups) and learning (h, N = 14, 8. Each replicate contains 12-14 flies) induced by Gr64f activation. For h, One-way ANOVA with Tukey’s multiple comparison test, ns. p>0.05; *** p<0.001. For j, *t* test, * p<0.05.

To ask if fox neurons function as a node in the neural pathway that connects sweet-sensing gustatory receptor neurons (GRNs) to PAM-DANs through FDA neurons, we performed a computational simulation^29^ with a leaky integrate-and-fire model based on the *Drosophila* whole brain connectome^19–21^. We simulated unilateral activation of labellar sweet-sensing GRNs at various frequencies and model synaptic weights. As expected, the number of PAM-DANs that responded gradually increased with increasing stimulation frequencies and model synaptic weights (Figure 5B, "fox intact" panel; Table S2). Silencing fox neurons almost completely abolished PAM-DAN firing at all conditions (Figure 5B, "fox silenced" panel; Table S2). Next, we tested silencing FDA neurons individually and in groups.

Silencing FDA Neuron-I, FDA Neuron-II, or all FDA neurons simultaneously significantly decreased the PAM-DANs response to GRN stimulation (Figure 5C; Table S1B). Interestingly, silencing CB0233 (an FDA Neuron-II that accounts for 7% of total fox synaptic output and 41% of fox synaptic output to FDA neurons) alone was sufficient to block PAM-DAN response (Figure 5C; Table S1B). Based on these simulations, we made two predictions: first, fox neurons should be functionally connected to PAM-DANs through their downstream ascending neurons; second, blocking fox neurons or their downstream ascending neurons should compromise appetitive learning performance.

To test the connection between fox neurons and PAM-DANs, we performed a functional connectivity assay using two-photon calcium imaging. Briefly, we optogenetically stimulated fox neurons while collecting calcium signals from PAM-DANs. We found that stimulating fox neurons activated PAM-DANs that innervated the MB γ4, γ5, β2, and β’2 compartments (Figure 5D), which play essential roles in appetitive olfactory learning^13,26^. To test if the connection between fox neurons and PAM-DANs can shape behaviors, we performed a neural epistasis experiment where we activated fox neurons while optogenetically blocking PAM-DANs. We found that blocking PAM-DANs did not affect increased food consumption caused by fox activation (Figure 5E) but completely abolished the learning performance generated by fox activation (Figure 5F), suggesting that fox neurons and PAM-DANs function sequentially in mediating learning.

To further test the connection between GRNs, fox, FDA neurons, and PAM-DANs, we activated neurons expressing sweet GRN Gr64f while blocking fox or CB0233 (FDA Neuron-II) neural activity. We found that blocking either fox or CB0233 neurons significantly suppressed the increased feeding phenotype and also compromised learning generated by Gr64f activation (Figures 3A, 4B, 5G, and 5H). These results collectively suggest that fox and FDA neurons are essential to convey the positive valence of sweet GRN activation to PAM-DANs for learning.

## Discussion

In this study, we identified a pair of neurons in *Drosophila* SEZ, termed fox neurons, as a hub linking sweet gustatory receptor neuron input to appetitive learning circuits. Our imaging and behavioral experiments support a working model where fox neurons assign and relay the valence of sweet GRN activity to PAM-DANs to facilitate learning, effectively bridging peripheral taste detection and central reinforcement pathways^12,13^ (Figure 5A).

This model is supported by several complementary lines of evidence: First, flies perceive the activation of fox neurons as rewarding. It both induces appetitive behavioral responses and serves as a positive reinforcer to generate appetitive learning (Figures 1B, E, F and 4A). Second, the increased food, water, and quinine consumption upon fox activation is constrained by food and water deprivation and the level of bitterness (Figures 1E, F, 2D), indicating that fox-induced appetitive drive remains regulated by homeostatic state rather than triggering a compulsive motor program. Third, we found fox activation can override the intrinsic neutral or aversive valence of otherwise unpalatable or bitter tastants, leading to consumption (Figures 2B-D). Conversely, silencing fox impairs sweet-GRN-driven increased consumption without compromising the recognition of taste identity (Figures 3C-E). Thus, fox activity confers a positive valence, not sensory detection, of tastants. Last, fox neurons relay the positive valence of tastants to the learning pathway through their downstream ascending neurons. Blocking either fox or these downstream neurons abolishes appetitive learning generated by sweet-GRN activation (Figures 4B and 5F, H). This requirement demonstrates that fox is essential for conveying reward signals to dopaminergic learning circuits.

Fox neurons implement a convergent–divergent motif in the gustatory circuit. Multiple peripheral sweet GRNs and GINs converge onto fox neurons, and fox outputs diverge to multiple targets (e.g., PAM-DANs, feeding motor centers). This is further supported by our computational simulations that showed blocking fox neurons and their ascending downstream neurons suppressed the global neuronal responses to gustatory neuron stimulation (Figure S6A and Table S3). This is a unique characteristic of fox neurons that is not shared by other SEZ cholinergic neurons with a similar bilateral morphology (termed ‘fox-like’ neurons, Figure S6B and Table S3). This “hourglass” structure ensures that sweet signals pass through a unified hedonic assignment node before distribution––a design that confers both consistency and flexibility. After this bottleneck, the reward signal is broadcast to separate downstream networks governing memory and feeding. For example, internal state signals may modulate fox neuron excitability at this bottleneck (e.g., through DSK neurons), thereby scaling the appetitive value of sweet tastes according to the animal’s needs. Such hourglass architectures are conserved across taxa: in mammals, for example, gustatory inputs from multiple peripheral fibers converge in the parabrachial nucleus before projecting to hypothalamic, amygdalar, and cortical targets involved in feeding and learning^30,31^. Such architectures––whether in insects or mammals––underlie the ability of organisms to distill complex sensory information into a unifying affective code that can drive coherent adaptive behaviors.

Our research also opens several important questions for future research. One particularly compelling direction is to characterize how fox and downstream neurons interact with pathways that assign negative valence. Another is to identify non-gustatory inputs to fox, e.g., to determine whether and how internal state indicators modulate fox activity.

Taken together, we propose that the fox circuitry represents a mechanism through which the positive valence of tastants is assigned and conveyed to the central brain. The role of fox neurons in *Drosophila* underscores the importance of specialized neural hubs for taste valence processing, providing insight into the neural basis of reward, motivation, and behavioral flexibility in the face of changing environments and needs.

## Acknowledgements

We thank Dr. Ulrike Heberlein for her support during early stages of this project and her feedback on the manuscript. We thank Yoonwoo Park and Sabina Knox for their contributions to the early stage of this project. We thank Drs. Jon-Michael Knapp, Yichun Shuai, Yoshi Aso, and Yufeng Pan for discussion and feedback. We thank Drs. Daniel Bushey, Glenn Turner, Mehrab Modi, Yoshi Aso, and Vasily Goncharov for technical support and advice. We are grateful to Dr. Key Ito, Yichun Shuai, Daniel Bushey, Glenn Turner, Yoshi Aso, Karla Kaun, Gabriella Sterne, Mark Wu, the HHMI Janelia Research Campus, and Bloomington *Drosophila* Stock Center (BDSC) for providing fly strains. We thank Dr. Philip Shiu for publicly sharing code for brain simulations and providing advice on implementation. We acknowledge the Princeton FlyWire team and members of the Murthy and Seung labs for development and maintenance of FlyWire (supported by BRAIN Initiative grant MH117815 to Murthy and Seung). This work was supported by UD-GUR, UD-UDRF, Delaware CTR program with a grant from the National Institute of General Medical Sciences (NIGMS, U54 GM104941), Delaware INBRE program with a grant from the NIGMS (P20 GM103446), and Maximizing Investigators’ Research Award to L.S. (NIGMS R35GM147504).

## Author contributions

L.S. conceived and supervised the project, conducted calcium imaging, positional preference, and associative learning experiments, and performed connectome analyses. P.C. and L.S. performed the neuroanatomy screen. M.I. generated the split-GAL4 drivers used in the neuroanatomy screen.

K.W.C. conducted calcium imaging experiments and performed connectome analyses and model simulations. T.S.D. performed immunostaining, confocal imaging, and feeding experiments. K.W.C., T.S.D., and L.S. performed data analyses and generated figures. L.S. wrote the manuscript with input from all authors.

## Materials and methods Key resources table

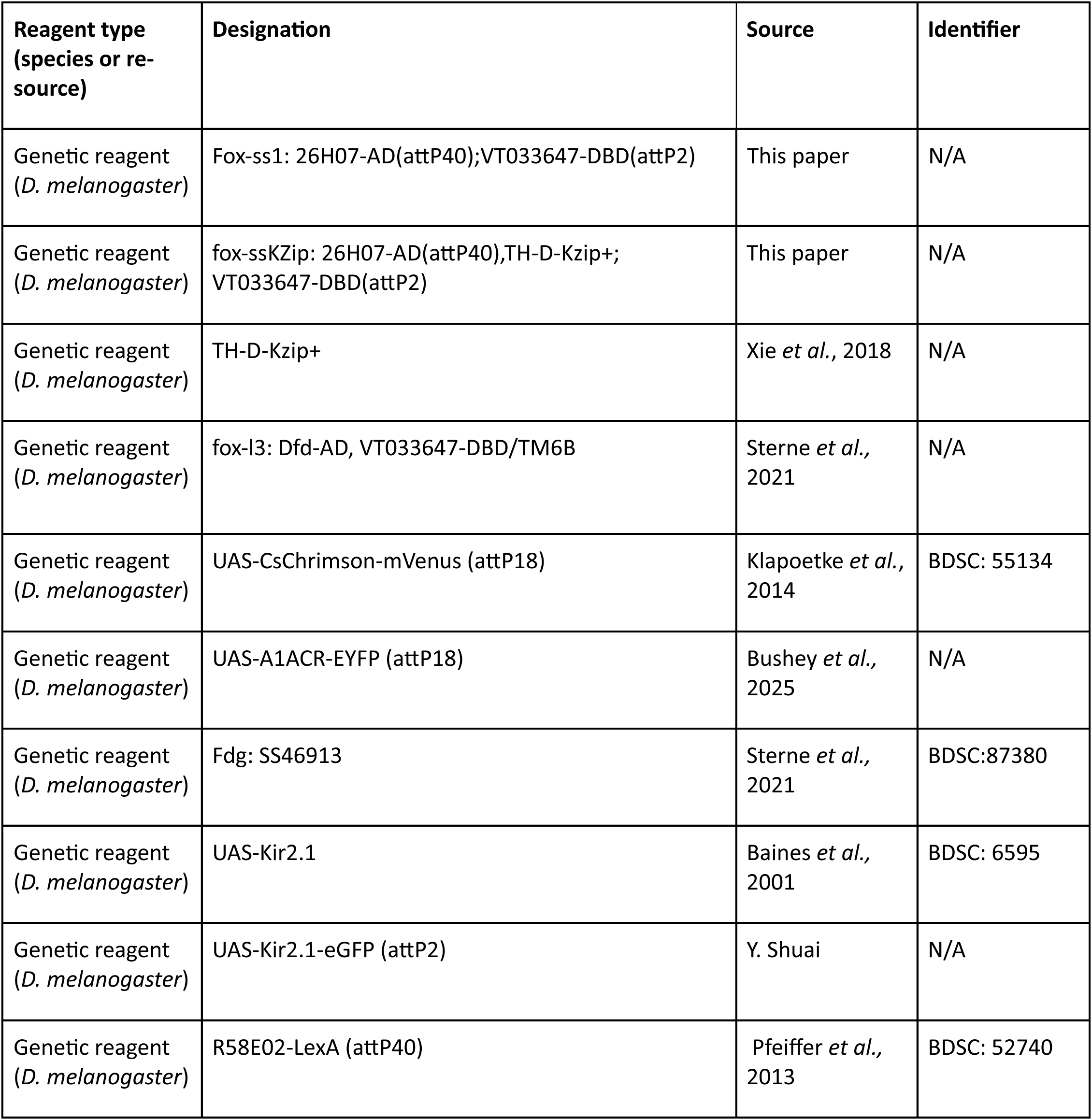
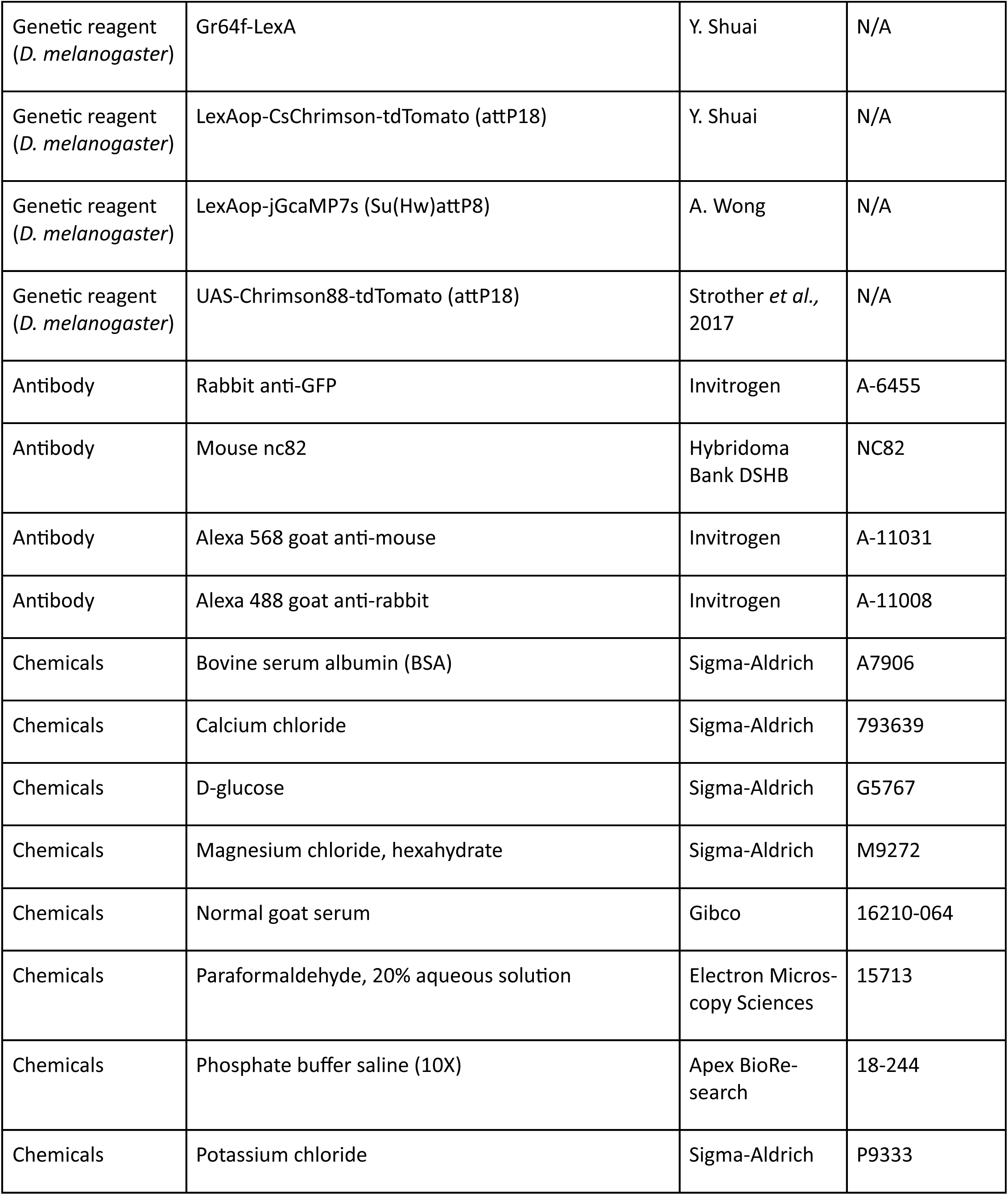
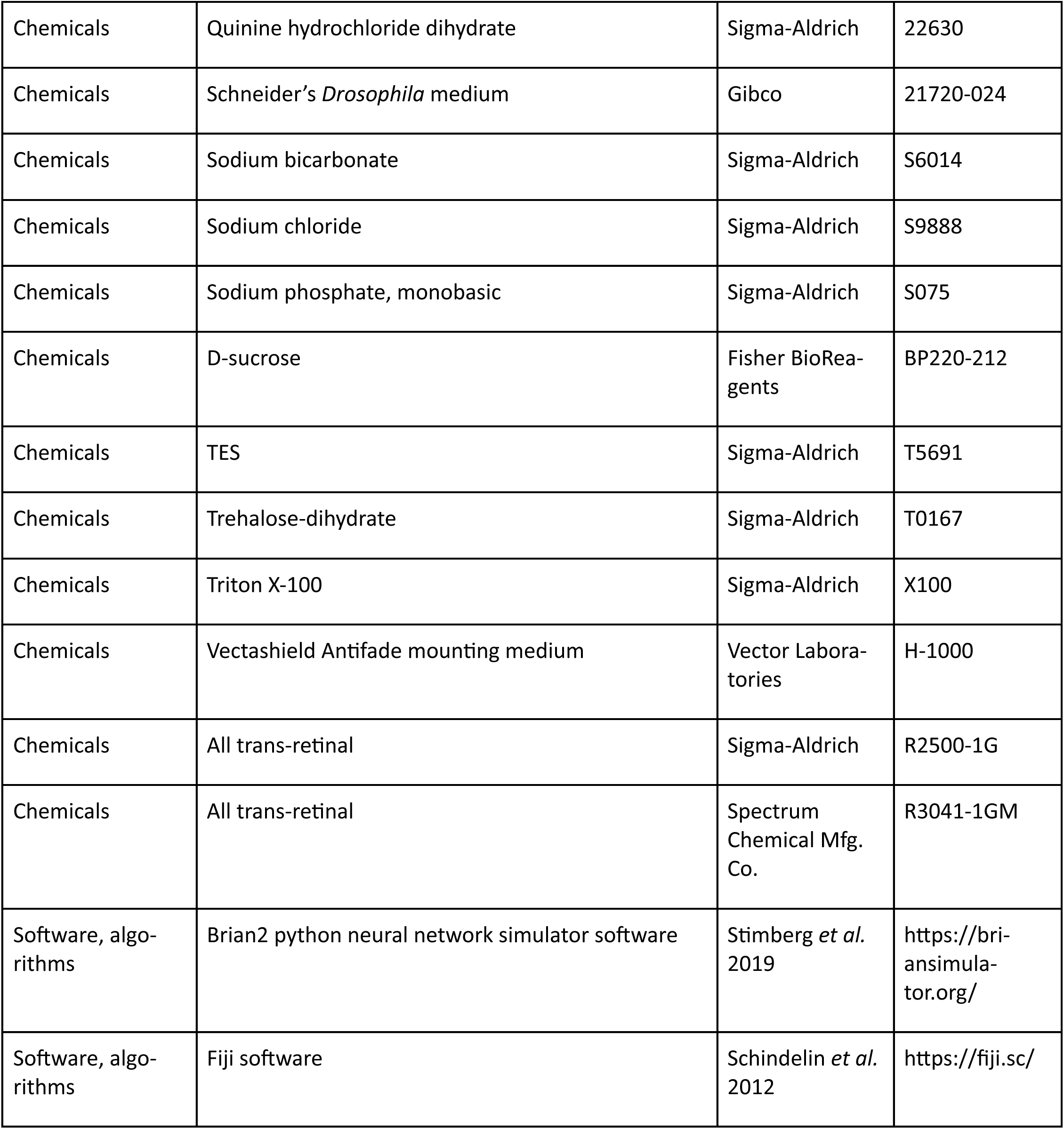
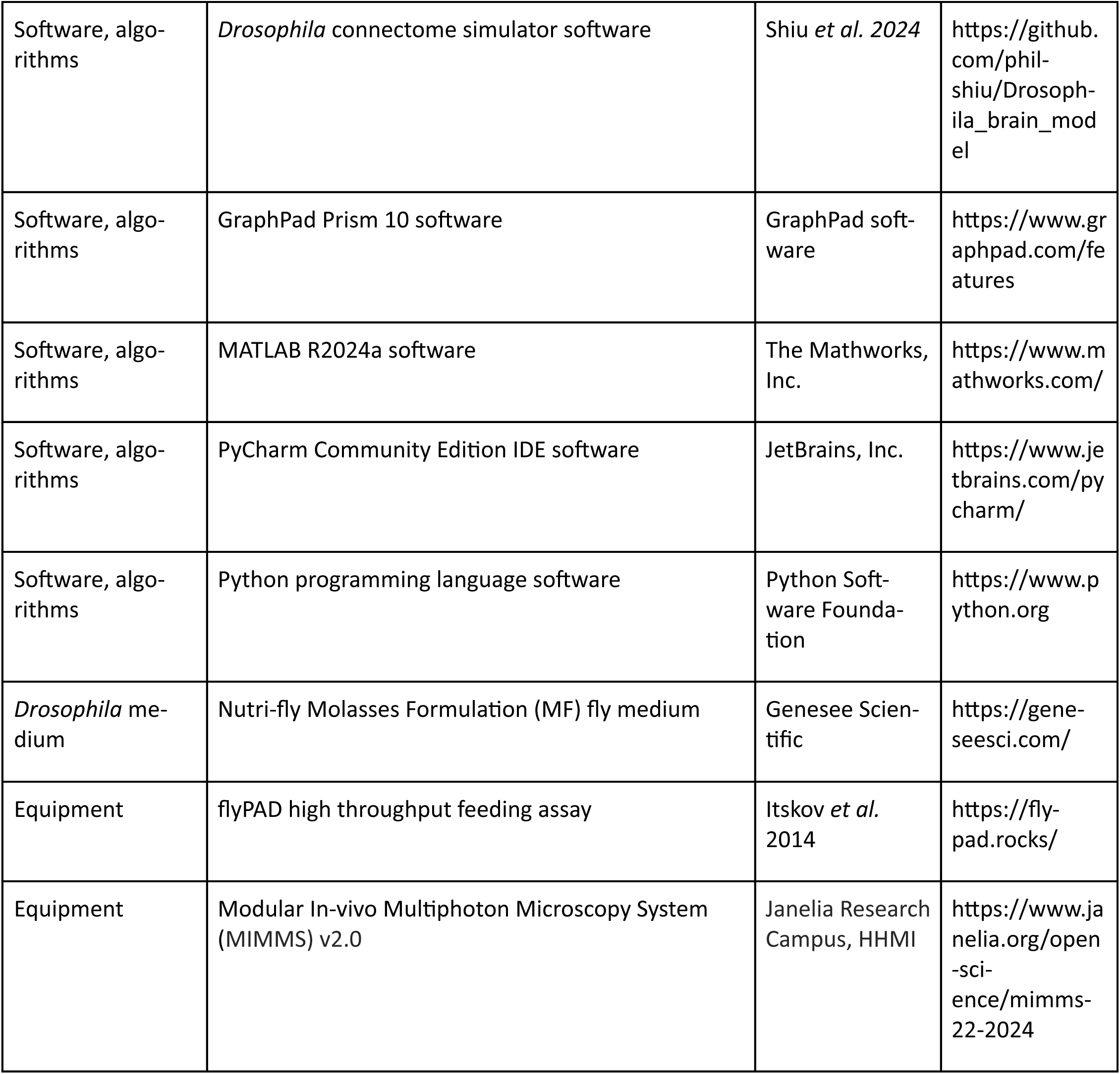

### Fly stocks and culture

*Drosophila melanogaster* stocks (listed in key resources table) were raised at 25°C and 50%-60% relative humidity on commercially prepared standard media (cornmeal/agar/molasses/yeast; Nutri-Fly MF) for anatomical and ‘Con-Ex’ feeding experiments. For optogenetic behavioral and *ex vivo* functional imaging experiments, flies were raised under constant darkness at 25°C and 50%-60% relative humidity on the same media supplemented with 0.2 mM all trans-retinal. Flies were collected 0-3 days post-eclosion on media containing 0.4 mM all trans-retinal. Flies for anatomical and behavioral assays were 4- to 7-days-old at the time of experiments; flies used for *ex vivo* functional connectivity calcium imaging were 2- to 5-days-old at the time of dissection.

### Behavioral assays

Positional preference assays were performed at 25°C, 50%-60% relative humidity in darkness. Female flies (3-7d old) were loaded into rectangular chambers, 8 flies per chamber. After a 2-min baseline period, one side of the chamber (selected at random) was illuminated by 0.5 Hz 1-second pulsed 625-nm red LED, at 40% intensity (100% intensity = 3.9 mW/cm^2) for 3 min. Following the illumination period, there was a 2-min recovery period. Experiments were recorded with infrared cameras (BlackFly USB3 camera, Teledyne vision solutions). The frequencies and intensities of LED stimulation were controlled using MATLAB (MathWorks, Inc.). Acquired video was analyzed with custom MATLAB scripts that detected the position of the flies, allowing for the calculation of a Preference Index as described in Shao *et al.* 2017.

Associative learning with optogenetic stimulation was conducted in a circular arena that couples optogenetic stimulation with an odor delivery system, as described in Aso *et al.,* 2014. All tests were performed at 25°C, 50%-60% relative humidity. The olfactory cues used were 4-methyl-cyclohexanol (1:1000) and 3-octanol (1:1000). Optogenetic stimulation used a 0.5 Hz 625-nm LED light at 40% or 10% intensities (100% intensity = 3.9 mW/cm^2). Flies were trained as groups of 12-14 individuals using a single training session consisting of 1-min exposure to air, followed by a 1-min exposure to odor A paired with neuronal activation/silencing, followed by a 1-min exposure to air, and finally a 1-min exposure to odor B. Memory was tested 1 min after training, during which the odors A and B were delivered to each pair of opposing quadrants. Experiments were recorded under IR illumination with a camera (Point Gray Research) located above the chamber and equipped with an IR long-pass filter. The subsequent video was analyzed with custom MATLAB scripts that detect the position of flies within the chamber.

To quantify 24-hour feeding behavior, we used an assay generally based on the ‘Con-Ex’ protocol described by Shell *et al.,* 2018. Dyed food was prepared via the addition of FD&C blue No. 1 dye to normal fly food with a concentration of 1% (wt/vol). A small volume of dyed food (100 μL) was pipetted into the underside of a 2.0 mL Eppendorf tube cap 30 min prior to adding flies. Two holes were poked near the top of the vials using a heated needle, allowing air/moisture circulation. Two 3- to 7-days-old mated female flies were transferred and allowed to feed *ad libitum* in the vials for 24 h at 25℃ (55% humidity). After feeding, vials were placed on dry ice to euthanize the flies, which were then transferred to a 1.7 mL Eppendorf tube. Fly carcasses were ground in 300 μL of distilled water and centrifuged. A portion of supernatant (100 μL) was transferred to a 96-well culture plate. Concurrently, the excreta in the feeding vial were dissolved in 500 μL of distilled water, vortexed for 5-10 s, and 100 μL of the sample transferred to the same 96-well culture plate. Using a plate reader, the absorbance values of the samples (λ = 620 nm) were measured and converted to concentrations via standard curves, then converted to usable mass for data analysis.

Acute feeding behavior was quantified by use of the FlyPad behavioral monitoring system (Itskov *et al.* 2014), at 25°C, 50% -60% relative humidity. Warmed liquified media was loaded into both probes in each FlyPAD chamber in darkness. Individual female flies (3-7 d old) were placed in each FlyPAD chamber. Flies were allowed to feed *ad libitum* for 50 min total. In the open-loop feeding and choice assay, the FlyPAD chambers were illuminated with 625-nm red LEDs to optogenetically stimulate channelrhodopsin-expressing neurons. In the closed-loop feeding assay, the FlyPAD chambers were only illuminated with red LED when flies fed on one of the two probes in each arena. The probe associated with illumination was alternated for the next set of experiments to mitigate potential probe-specific bias. Media containing 7% yeast, 160 mM sucrose, and 0.7% agar was used for open-loop feeding assay. Media containing either 0.7% agar (neutral taste), 100 mM sucrose (sweet taste), or 0.5 mM quinine (bitter taste) were used for open-loop choice assays and closed-loop feeding assays. Analysis of feeding behavior used custom MATLAB scripts that extracted the total number of sips during the feeding period.

### Immunostaining and confocal imaging

Immunohistochemical staining and imaging were performed as previously described (Shao *et al.,* 2019). Briefly, brains were dissected in cold Schneider’s Insect (S2) medium, fixed in 2% paraformaldehyde in S2 at Room Temperature (RT) under nutation for 55-65 min in covered Eppendorf tubes, washed 3-4 times with PBT0.5 (phosphate buffered saline, PBS, containing 0.5% Triton X-100) at RT, blocked with PAT3 (1xPBS, 1g/100mL BSA, 0.5% Triton X-100) for 1 hour at RT, incubated with primary antibody (1:500 rabbit anti-GFP, 1:50 mouse anti-nc82 in PAT3) at 4°C overnight, washed 3-4 times in PBT0.5, incubated with secondary antibody (1:500 Alexa 488 antirabbit, 1:500 Alexa 568 anti-mouse in PAT3) at 4°C overnight, washed with PBT0.5 and PBS, and mounted in Vectashield. The brains were then imaged on a confocal microscope (Zeiss CellDiscoverer 7, using Zen Blue software).

### *Ex vivo* calcium imaging

All functional imaging was performed on 2- to 5-days-old female flies. Whole brains were dissected in a Sylgard silicone elastomer (Dow Inc.) coated dish, immersed in artificial fly hemolymph (103 mM NaCl, 3mM KCl, 2 mM CaCl_2_, 4 mM MgCl_2_·6 H_2_O, 26 mM NaHCO_3_, 1 mM NaH_2_PO_4_, 8 mM trehalose, 10 mM glucose, 5 mM TES, bubbled with 95% O_2_/5% CO_2_). Brains were placed on a poly-Lysine coated coverslip and imaged with a resonant-scanning 2-photon microscope (MIMMS v2.0, Janelia Research Campus), using a laser with a 920 nm excitation wavelength (Toptica FemtoFiber Ultra 920) and an excitation power of 19-33 mW (2.7-4.3 mW/mm^2^ intensity). Emitted fluorescence was detected with GaAsP photomultiplier tubes (Hamamatsu) controlled via a Janelia PMT Controller. Images were acquired with an Olympus × 40 water-dipping, 0.8 numerical aperture objective at 512 pixels × 512 pixels resolution. Optogenetic stimulation of CsChrimson was provided by a 660-nm red light LED diode (ThorLabs M660L4), fed into a digital mirror device (Texas Instruments DLP Lightcrafter 4500) and projected through the objective. Photostimulation consisted of 4 x 1 s pulses, separated by a 24 s inter-pulse interval, with a stimulation power of 20-21 𝜇W (2.8-3.0 μW/mm^2^ intensity). Volume stacks were continuously imaged at 2x magnification, and consisted of 45 slices, with an inter-slice interval of 2 𝜇m, allowing all the PAM-DANs compartments to be imaged. Stack acquisition began 10s prior to the 1st LED pulse and stopped 24s after the termination of the final pulse. After optogenetic stimulation, brains were imaged for ROI definition of mushroom body compartment anatomy using the same imaging stack dimensions as above, using both 920 nm, and 1050 nm excitation (Toptica FemtoFiber Ultra 1050). Then power intensity for imaging at the 920- and 1050-nm excitation was 3.2-3.4 and 3.1-3.2 mW/mm^2^ respectively.

Regions of interest (ROI)s for each brain were drawn using custom MATLAB software and the 1050 nm anatomy images described above. The average pixel intensity for each ROI (𝑓𝑙𝑢𝑜𝑟_𝑎𝑣#_) was calculated, as was its average baseline intensity (𝑓𝑙𝑢𝑜𝑟_𝑏𝑠𝑒𝑙𝑛_), defined as the averaged intensity for the 5 volumes (approximately 1.5 s) acquired immediately prior to each optogenetic pulse onset. Average background intensity (𝑓𝑙𝑢𝑜𝑟_𝑏𝑐𝑘#𝑟_) for each volume was calculated from a ROI located in non-brain-tissue containing space. For each PAM compartment, ΔF/F was calculated using the formula:

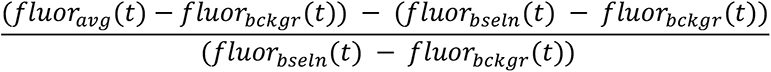

## Connectome simulation

Digital experiments were performed using a Python-based *Drosophila melanogaster* neural simulation, described previously (Shiu *et al.* 2024), implemented using the open-source Brian2 library (Stimberg *et al.* 2019). The default model parameters described by Shiu *et al.* 2024 were used: 1000 ms trial duration, 30 trial repetitions, −52 mV resting potential, −52 mV reset potential after a spike, −45 mV spiking threshold, 2.2 ms refractory period, 5 ms tau for synapse decay, 1.8 ms time delay from spike to change in membrane potential. The free parameter model synaptic weight (W_syn_) was adjusted to achieve differing levels of propagation of GRN activation throughout the connectome. Labellar sugar gustatory neurons (GRNs, Shiu *et al.* 2024), second- and third-order gustatory interneurons (GINs, Shiu *et al.* 2023), and fox neurons were stimulated using Poisson distributed input at stimulation rates from 10-200 Hz. A downstream neuron was considered activated if its average firing rate was non-zero during stimulation of any of the simulated experiments. Experiments were executed in Python, using the PyCharm Community Edition IDE, and the output (mean spiking rate tables of activated connectome neurons) were used for further analysis and visualization using custom MATLAB scripts (The MathWorks, Inc).

## Statistical analysis

Statistical analysis was conducted using GraphPad Prism. Unless otherwise stated, unpaired *t* tests or one-way ANOVA tests were performed to compare between genotypes and conditions, with Tukey post-hoc testing (α = 0.05). All data are expressed as means ± SEM.

