## Supplemental Figures and Tables for "From perception to valence: a pair of interneurons that assign positive valence to sweet sensation in *Drosophila*"

### Supplementary Figures

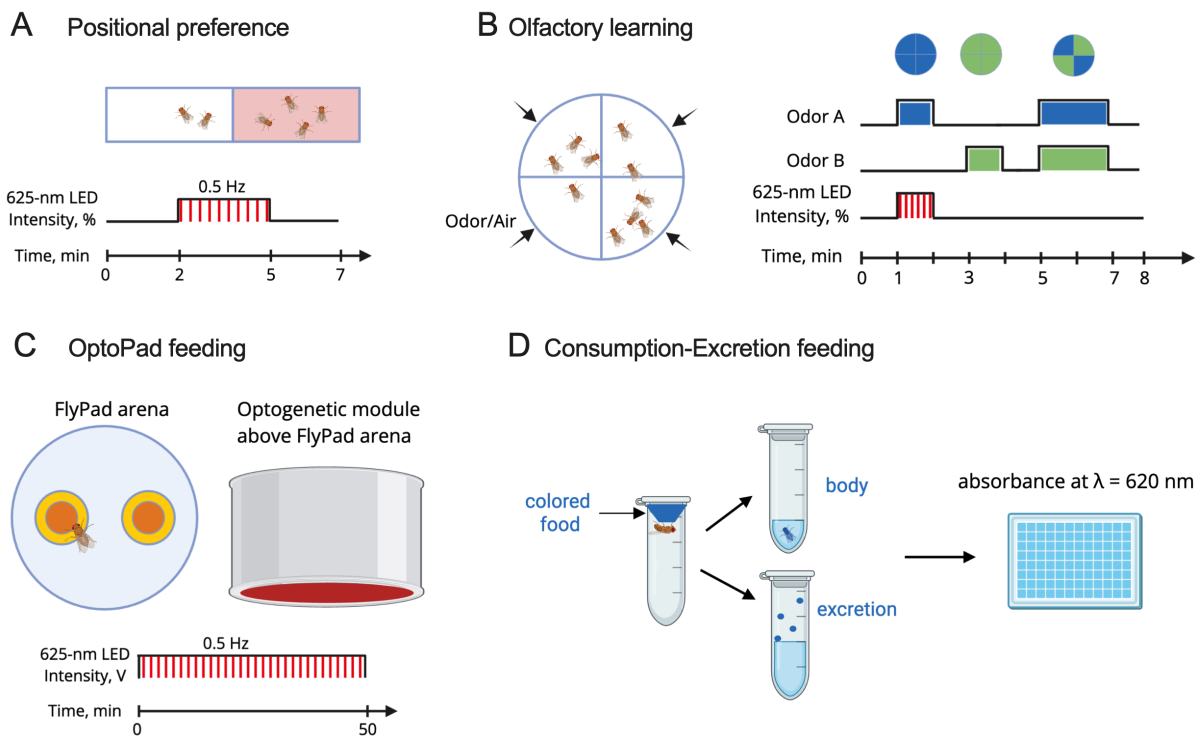


Figure S1. Schematics for behavior assays

A. Positional preference assay: Flies were first loaded into a rectangular arena. Following a 2-minute acclimation period, half of the arena was illuminated with a 625-nm LED for 3 minutes. A subsequent 2-minute recovery period was provided after illumination.

B. Olfactory learning assay: Flies were introduced into a circular arena that was subdivided into four quadrants to enable localized delivery of odors or air. After a 1-minute adaptation period, the arena was illuminated with a 0.5 Hz 625-nm LED while odor A was administered for 1 minute. This was followed by a 1-minute airflow to clear residual odor. Next, odor B was delivered for 1 minute without LED illumination. Finally, after another clearing period, both odors A and B were simultaneously presented from opposite quadrants for 2 minutes to enable a choice.

C. OptoPad feeding assay: This is a FlyPad assay with optogenetic function. A single fly was loaded into a FlyPad arena containing solid food placed at the center of metal rings. Feeding events were recorded when the fly, by contacting the food, completed an electrical circuit. During the open-loop assay, the arena was continuously illuminated by the optogenetic module from above with a 625-nm LED for 50 minutes. Control flies are allowed to feed in the dark, without optogenetic illumination. During the closed-loop assay, the arena was illuminated for 2 seconds by optogenetic module only when the fly is feeding on one of the probes. The probe associated with activation was alternated for each trial to counterbalance potential intrinsic preference for one probe.

D. Consumption-Excretion (Con-Ex) feeding assay: Flies were maintained on colored food provided in a feeding tube over a 24-hour period. The amount of food consumed was quantified, based on the dye absorbance by recovering and measuring the blue dye from both the homogenized flies and the excretion residues washed from the walls of the feeding tube.

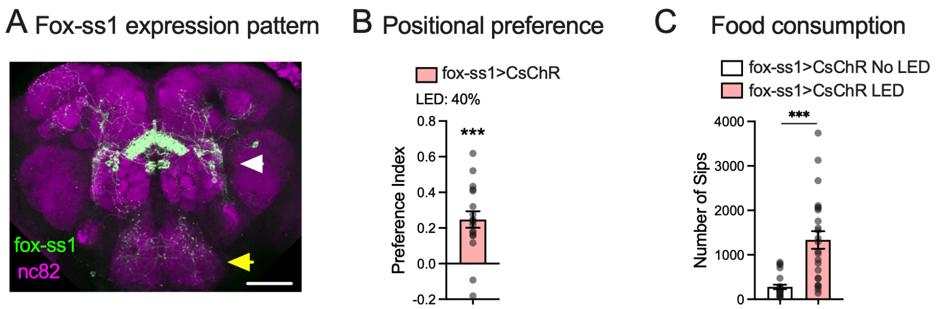


Figure S2. Expression pattern and phenotypes of original fox driver (fox-ss1)

A. Immunostaining of CsChrimson-mVenus driven by fox-ss1 driver lines that labels fox neurons (yellow arrow) and PPM3 dopaminergic neurons (white arrow).

B. Flies preferred the illuminated side in the positional preference assay when their fox neurons were optogenetically activated (fox-ss1>CsChrimson). N = 18. Each replicate contains 6-8 flies. One-sample *t* test (hypothetical value = 0), *** $p<0.001$.

C. Activating fox neurons with fox-ss1 driver increased consumption of food in fed flies. N = 24 flies. *t* test (no LED and LED groups of the same genotype), *** $p<0.001$.

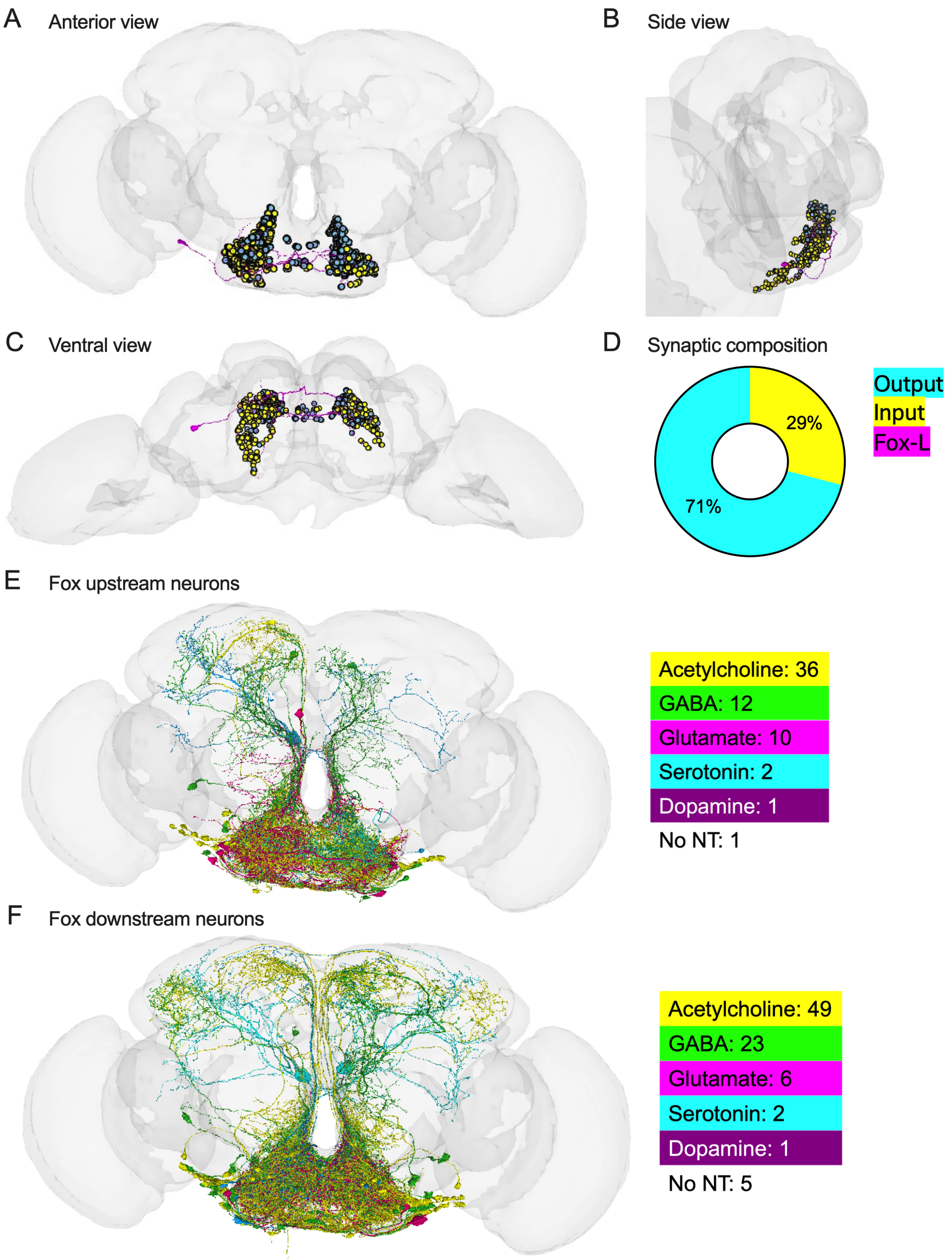


Figure S3. Fox neurons in FAFB.

A-C. Anterior, side, and ventral views of the left-hemisphere fox neuron in FAFB. Yellow dots indicate synaptic inputs; blue dots indicate synaptic outputs.

D. Synaptic composition of the left-hemisphere fox neuron.

E-F. The upstream and downstream synaptic partners of the left-hemisphere fox neuron.

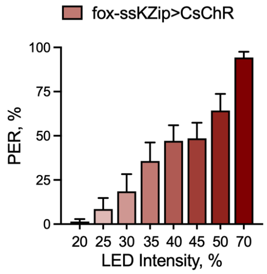


Figure S4. Activating fox neurons induces the proboscis extension reflex

Activating fox neurons (fox-ssKZip>CsChrimson) induced proboscis extension in a dosage-dependent manner. N = 14 flies.

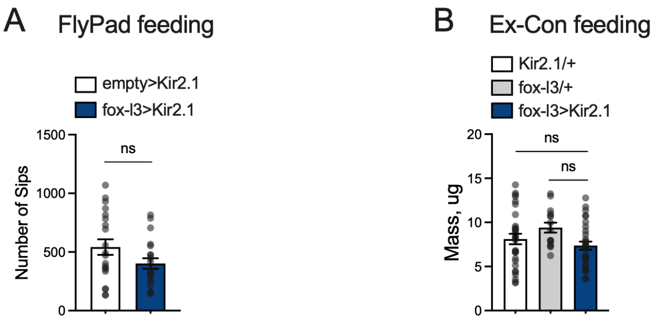


Figure S5. Blocking fox neurons did not affect feeding in fed flies.

A. Constitutively blocking fox neurons with Kir2.1 (fox-l3>Kir2.1) did not affect food consumption in the FlyPad feeding assay compared with genetic control (empty>Kir2.1). N=21 flies for empty>Kir2.1; N=24 flies for fox-l3>Kir2.1. *t* test, ns. p>0.05.

B. Constitutively blocking fox neurons with Kir2.1 (fox-l3>Kir2.1) did not affect 24-hour food consumption compared with genetic controls (Kir2.1/+ and fox-l3/+). N = 30 flies for Kir2.1/+ and fox-l3>Kir2.1; N = 15 flies for fox-l3/+. One-way ANOVA with Tukey's multiple comparison test, ns. P>0.05.

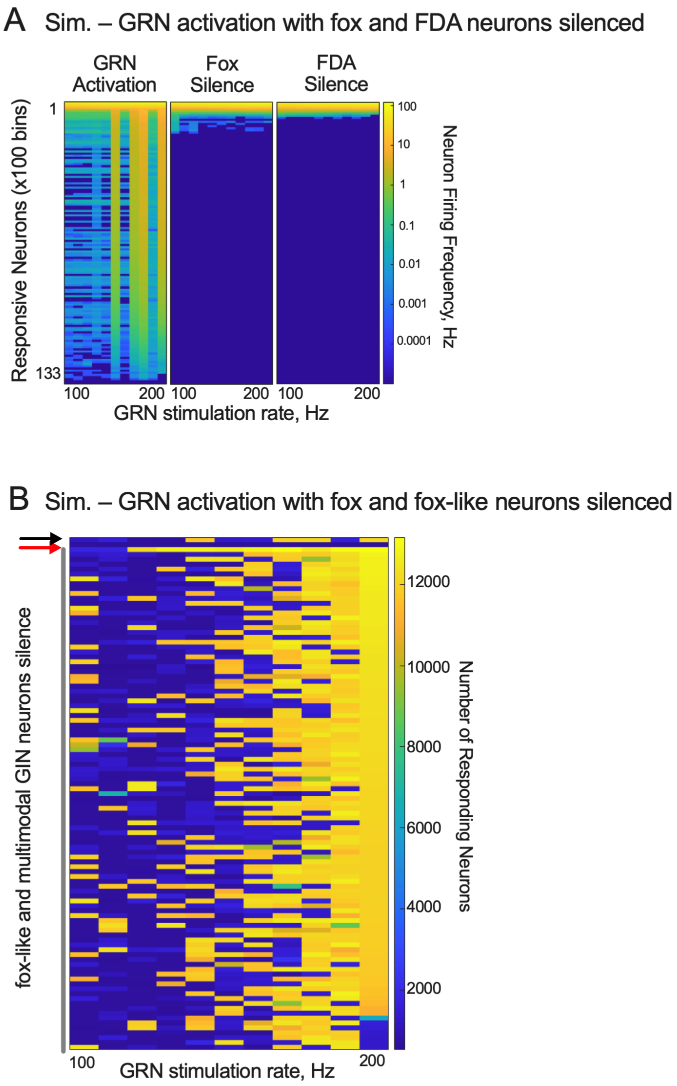


Figure S6. Fox neurons facilitate relaying the positive valence of tastants to the central brain.

A. Simulation of stimulating labellar sweet GRNs without (left panel) and with fox neurons (middle panel) and FDAs (right panel) blocked in computational simulation (model synaptic weight = 0.39), shown as the firing frequency of responsive neurons (averaged 100 neurons/bin).

B. Simulation of blocking the fox-like neurons and multimodal GINs while GRNs were stimulated in computational simulation (model synaptic weight = 0.39). The black arrow indicates the results of GRN activation; the red arrow indicates the results of silencing fox. The grey line indicates the results of silencing fox-like neurons and multimodal GINs (these categories overlap). The heatmap represents the number of responsive neurons in each simulation trial.

**Supplementary Tables**

Table S1A. Simulated number of responding neurons in the connectome to sugar GRN stimulation during silencing of FDA neuronal pairs or sets.

| Silenced neuron(s) | Cell type | 20 (Hz) | 40 (Hz) | 80 (Hz) | 100 (Hz) | 110 (Hz) | 120 (Hz) | 130 (Hz) | 140 (Hz) | 150 (Hz) | 160 (Hz) | 170 (Hz) | 180 (Hz) | 190 (Hz) | 200 (Hz) |
|---|---|---|---|---|---|---|---|---|---|---|---|---|---|---|---|
| No silencing | | 644 | 659 | 1498 | 1185 | 1606 | 635 | 658 | 11720 | 1321 | 1431 | 11946 | 12043 | 1616 | 12169 |
| GNG.SMP.3 / GNG.SMP.8 | CB0233 | 1487 | 656 | 613 | 656 | 595 | 620 | 599 | 595 | 602 | 618 | 593 | 574 | 610 | 577 |
| GNG.SMP.10 / PRW.SMP.4 | CB0546 | 690 | 686 | 630 | 11056 | 11972 | 645 | 624 | 637 | 650 | 658 | 598 | 1294 | 11825 | 635 |
| SLP.GNG.3 / SLP.GNG.1 | DNp62 | 645 | 1266 | 638 | 11848 | 629 | 638 | 11328 | 2467 | 11674 | 1314 | 12022 | 11954 | 1691 | 12053 |
| PRW.SMP.16 / SMP.163 | CB0272 | 697 | 667 | 673 | 633 | 627 | 629 | 12467 | 11566 | 1307 | 623 | 1455 | 11943 | 11269 | 12201 |
| GNG.79 / GNG.128 | CB0337 | 11740 | 747 | 638 | 649 | 654 | 11825 | 677 | 1532 | 11821 | 12000 | 12353 | 11009 | 12278 | 11959 |
| GNG.SMP.17 / FLA.SMP.61 | CB1514 | 9789 | 756 | 638 | 1319 | 630 | 662 | 621 | 1124 | 11436 | 12454 | 1636 | 12287 | 11793 | 1295 |
| FLA.SMP.58 / FLA.SMP.61 | CB1514 | 12149 | 695 | 645 | 1186 | 676 | 1181 | 622 | 1346 | 1331 | 643 | 1465 | 11898 | 12672 | 11930 |
| GNG.SMP.18 / GNG.SMP.24 | CB3199 | 1266 | 707 | 656 | 625 | 633 | 11821 | 1257 | 627 | 1570 | 654 | 12239 | 11454 | 11947 | 12326 |
| SMP.1281 / SMP.1493 | CB1025 | 1719 | 655 | 657 | 11669 | 1260 | 630 | 1150 | 646 | 1147 | 1208 | 12504 | 11824 | 11003 | 12708 |
| GNG.SMP.19 / SMP.2023 | CB3470 | 1355 | 689 | 1167 | 667 | 627 | 644 | 660 | 12409 | 1346 | 1272 | 11841 | 11402 | 11946 | 12191 |
| SMP.1086 / SMP.1499 | CB3573 | 11386 | 660 | 646 | 1286 | 627 | 633 | 1265 | 12407 | 1058 | 1261 | 12329 | 1605 | 11934 | 12865 |
| FDA-I | | 655 | 1331 | 673 | 654 | 621 | 12371 | 620 | 634 | 626 | 617 | 1407 | 612 | 11792 | 1194 |
| FDA-II | | 1320 | 625 | 606 | 600 | 603 | 632 | 632 | 589 | 597 | 592 | 592 | 594 | 589 | 576 |
| FDA-All | | 634 | 1355 | 621 | 585 | 599 | 594 | 603 | 599 | 594 | 587 | 614 | 593 | 583 | 592 |

Table S1B. Simulated number of responding PAM-DANs to sugar GRN stimulation during silencing of FDA neuronal pairs or sets.

| Silenced neuron(s) | Cell type | 20 (Hz) | 40 (Hz) | 80 (Hz) | 100 (Hz) | 110 (Hz) | 120 (Hz) | 130 (Hz) | 140 (Hz) | 150 (Hz) | 160 (Hz) | 170 (Hz) | 180 (Hz) | 190 (Hz) | 200 (Hz) |
|---|---|---|---|---|---|---|---|---|---|---|---|---|---|---|---|
| No silencing | | 0 | 0 | 8 | 6 | 8 | 0 | 0 | 200 | 7 | 11 | 201 | 203 | 10 | 201 |
| GNG.SMP.3 / GNG.SMP.8 | CB0233 | 11 | 0 | 0 | 0 | 0 | 0 | 0 | 0 | 0 | 0 | 0 | 0 | 0 | 0 |
| GNG.SMP.10 / PRW.SMP.4 | CB0546 | 0 | 0 | 0 | 0 | 202 | 0 | 0 | 200 | 7 | 0 | 7 | 199 | 196 | 205 |
| SLP.GNG.3 / SLP.GNG.1 | DNp62 | 0 | 7 | 0 | 201 | 0 | 0 | 198 | 10 | 199 | 8 | 202 | 201 | 9 | 202 |
| PRW.SMP.16 / SMP.163 | CB0272 | 0 | 0 | 0 | 0 | 0 | 0 | 201 | 200 | 7 | 0 | 7 | 199 | 196 | 205 |
| GNG.79 / GNG.128 | CB0337 | 200 | 0 | 0 | 0 | 0 | 201 | 0 | 8 | 200 | 202 | 203 | 196 | 200 | 200 |
| GNG.SMP.17 / FLA.SMP.61 | CB1514 | 173 | 0 | 0 | 10 | 0 | 0 | 0 | 7 | 196 | 200 | 11 | 202 | 201 | 7 |
| FLA.SMP.58 / FLA.SMP.61 | CB1514 | 200 | 0 | 0 | 6 | 0 | 7 | 0 | 7 | 8 | 0 | 11 | 200 | 200 | 202 |
| GNG.SMP.18 / GNG.SMP.24 | CB3199 | 6 | 0 | 0 | 0 | 0 | 201 | 9 | 0 | 7 | 0 | 201 | 199 | 201 | 200 |
| SMP.1281 / SMP.1493 | CB1025 | 7 | 0 | 0 | 197 | 9 | 0 | 6 | 0 | 6 | 8 | 203 | 199 | 194 | 205 |
| GNG.SMP.19 / SMP.2023 | CB3470 | 8 | 0 | 6 | 0 | 0 | 0 | 0 | 200 | 9 | 7 | 199 | 199 | 199 | 202 |
| SMP.1086 / SMP.1499 | CB3573 | 199 | 0 | 0 | 9 | 0 | 0 | 10 | 200 | 3 | 8 | 200 | 9 | 200 | 202 |
| FDA-I | | 0 | 7 | 0 | 0 | 0 | 200 | 0 | 0 | 0 | 0 | 7 | 0 | 200 | 7 |
| FDA-II | | 10 | 0 | 0 | 0 | 0 | 0 | 0 | 0 | 0 | 0 | 0 | 0 | 0 | 0 |
| FDA-All | | 0 | 9 | 0 | 0 | 0 | 0 | 0 | 0 | 0 | 0 | 0 | 0 | 0 | 0 |

Table S2. Simulated number of responding PAM-DANs to sugar GRN stimulation with and without silencing Fox, while varying simulation free parameter – synaptic weights (Wsyn)

| Condition | Wsyn | 10 (Hz) | 20 (Hz) | 30 (Hz) | 40 (Hz) | 50 (Hz) | 60 (Hz) | 70 (Hz) | 80 (Hz) | 90 (Hz) | 100 (Hz) | 110 (Hz) | 120 (Hz) | 130 (Hz) | 140 (Hz) | 150 (Hz) | 160 (Hz) | 170 (Hz) | 180 (Hz) | 190 (Hz) | 200 (Hz) |
|---|---|---|---|---|---|---|---|---|---|---|---|---|---|---|---|---|---|---|---|---|---|
| Fox intact | 0.37 | 0 | 0 | 0 | 0 | 0 | 0 | 0 | 0 | 0 | 0 | 0 | 0 | 0 | 0 | 0 | 191 | 0 | 5 | 8 | 189 |
| Fox intact | 0.38 | 0 | 0 | 0 | 0 | 0 | 0 | 0 | 0 | 0 | 0 | 0 | 0 | 7 | 8 | 0 | 0 | 8 | 8 | 6 | 196 |
| Fox intact | 0.382 | 0 | 0 | 0 | 0 | 0 | 0 | 0 | 0 | 0 | 0 | 0 | 0 | 0 | 0 | 0 | 8 | 198 | 7 | 6 | 186 |
| Fox intact | 0.386 | 0 | 0 | 0 | 0 | 0 | 0 | 0 | 0 | 0 | 0 | 0 | 0 | 0 | 0 | 0 | 0 | 201 | 201 | 7 | 201 |
| Fox intact | 0.39 | 0 | 0 | 0 | 0 | 0 | 0 | 0 | 8 | 0 | 6 | 8 | 0 | 0 | 200 | 7 | 11 | 201 | 203 | 10 | 201 |
| Fox silenced | 0.37 | 0 | 0 | 0 | 0 | 0 | 0 | 0 | 0 | 0 | 0 | 0 | 0 | 0 | 0 | 0 | 0 | 0 | 0 | 0 | 0 |
| Fox silenced | 0.38 | 0 | 0 | 0 | 0 | 0 | 0 | 0 | 0 | 0 | 0 | 0 | 0 | 0 | 0 | 0 | 0 | 0 | 0 | 0 | 0 |
| Fox silenced | 0.382 | 0 | 0 | 0 | 0 | 0 | 0 | 0 | 0 | 0 | 0 | 0 | 0 | 0 | 0 | 0 | 0 | 0 | 0 | 0 | 0 |
| Fox silenced | 0.386 | 0 | 6 | 0 | 0 | 0 | 0 | 0 | 0 | 0 | 0 | 0 | 0 | 0 | 0 | 0 | 0 | 0 | 0 | 0 | 0 |
| Fox silenced | 0.39 | 0 | 0 | 0 | 0 | 0 | 0 | 0 | 0 | 0 | 0 | 0 | 0 | 0 | 0 | 0 | 0 | 0 | 0 | 0 | 0 |

Table S3A. Simulated number of responding neurons in the connectome to sugar GRN stimulation during silencing of select neuronal pairs or single cells.

| Experiment | Silenced neuron(s) | 100 (Hz) | 110 (Hz) | 120 (Hz) | 130 (Hz) | 140 (Hz) | 150 (Hz) | 160 (Hz) | 170 (Hz) | 180 (Hz) | 190 (Hz) | 200 (Hz) |
|---|---|---|---|---|---|---|---|---|---|---|---|---|
| Control | No silencing | 1185 | 1606 | 635 | 658 | 11720 | 1321 | 1431 | 11946 | 12043 | 1616 | 12169 |
| Fox silence | CB0525 (Fox) | 663 | 603 | 639 | 618 | 617 | 592 | 622 | 622 | 608 | 619 | 646 |
| FDA silence | CB0233 | 656 | 595 | 620 | 599 | 595 | 602 | 618 | 593 | 574 | 610 | 577 |
| FDA silence | CB0546 | 11056 | 11972 | 645 | 624 | 637 | 650 | 658 | 598 | 1294 | 11825 | 635 |
| FDA silence | DNp62 | 11848 | 629 | 638 | 11328 | 2467 | 11674 | 1314 | 12022 | 11954 | 1691 | 12053 |
| FDA silence | CB0272 | 633 | 627 | 629 | 12467 | 11566 | 1307 | 623 | 1455 | 11943 | 11269 | 12201 |
| FDA silence | CB0337 | 649 | 654 | 11825 | 677 | 1532 | 11821 | 12000 | 12353 | 11009 | 12278 | 11959 |
| FDA silence | CB1514 (1) | 1319 | 630 | 662 | 621 | 1124 | 11436 | 12454 | 1636 | 12287 | 11793 | 1295 |
| FDA silence | CB1514 (2) | 1186 | 676 | 1181 | 622 | 1346 | 1331 | 643 | 1465 | 11898 | 12672 | 11930 |
| FDA silence | CB3199 | 625 | 633 | 11821 | 1257 | 627 | 1570 | 654 | 12239 | 11454 | 11947 | 12326 |
| FDA silence | CB1025 | 11669 | 1260 | 630 | 1150 | 646 | 1147 | 1208 | 12504 | 11824 | 11003 | 12708 |
| FDA silence | CB3470 | 667 | 627 | 644 | 660 | 12409 | 1346 | 1272 | 11841 | 11402 | 11946 | 12191 |
| FDA silence | CB3573 | 1286 | 627 | 633 | 1265 | 12407 | 1058 | 1261 | 12329 | 1605 | 11934 | 12865 |
| FDA silence | FDA-I | 654 | 621 | 12371 | 620 | 634 | 626 | 617 | 1407 | 612 | 11792 | 1194 |
| FDA silence | FDA-II | 617 | 603 | 586 | 632 | 613 | 585 | 598 | 601 | 571 | 589 | 582 |
| FDA silence | FDA-All | 585 | 599 | 594 | 603 | 599 | 594 | 587 | 614 | 593 | 583 | 592 |
| Foxlike silence | CB0152 | 617 | 633 | 626 | 10674 | 1208 | 1087 | 1098 | 9903 | 12002 | 12377 | 12054 |
| Foxlike silence | CB0117 | 1287 | 1240 | 625 | 619 | 622 | 1126 | 11744 | 12428 | 11839 | 12052 | 11677 |
| Foxlike silence | CB3793 | 644 | 1524 | 620 | 624 | 631 | 1354 | 1263 | 11687 | 12460 | 11911 | 1359 |
| Foxlike silence | CB0170 | 635 | 639 | 12470 | 1329 | 1431 | 1429 | 11588 | 11929 | 1276 | 1263 | 12070 |
| Foxlike silence | CB0251 | 620 | 632 | 629 | 633 | 1239 | 625 | 1256 | 642 | 11935 | 12389 | 12093 |
| Foxlike silence | CB0616 | 9462 | 1294 | 635 | 1313 | 11741 | 1246 | 1287 | 1185 | 11957 | 12319 | 12242 |
| Foxlike silence | CB0560 | 11112 | 645 | 632 | 635 | 627 | 1191 | 10776 | 10583 | 11042 | 12604 | 12370 |
| Foxlike silence | CB0553 | 528 | 552 | 521 | 1218 | 549 | 11628 | 1572 | 1290 | 12349 | 12065 | 12415 |
| Foxlike silence | CB0880 | 675 | 635 | 1472 | 628 | 11587 | 1188 | 1074 | 11523 | 12458 | 11856 | 12428 |

| | | | | | | | | | | | | |
|---|---|---|---|---|---|---|---|---|---|---|---|---|
| Foxlike silence | CB0848 | 1232 | 1335 | 641 | 1296 | 11989 | 630 | 12473 | 11659 | 11800 | 11981 | 12209 |
| Foxlike silence | CB0421 | 617 | 629 | 12198 | 665 | 639 | 1243 | 11640 | 10218 | 12338 | 11883 | 12495 |
| Foxlike silence | CB0247 | 1757 | 1779 | 12271 | 12831 | 12859 | 12690 | 12899 | 13125 | 12537 | 13028 | 13170 |
| Foxlike silence | CB0617 | 12446 | 1632 | 671 | 1318 | 12533 | 11661 | 785 | 947 | 12372 | 12119 | 12700 |
| Foxlike silence | CB0473 | 644 | 623 | 644 | 11445 | 607 | 11467 | 1384 | 12524 | 11871 | 12008 | 12323 |
| Foxlike silence | CB0331 | 625 | 643 | 11710 | 1334 | 1079 | 1275 | 12427 | 11982 | 12153 | 12686 | 12005 |
| Foxlike silence | CB0287 | 1279 | 1502 | 621 | 643 | 11852 | 1365 | 12317 | 1106 | 11807 | 12694 | 11897 |
| Foxlike silence | CB0791 | 662 | 657 | 649 | 669 | 823 | 11692 | 11759 | 11761 | 11844 | 12193 | 12275 |
| Foxlike silence | CB0768 | 11762 | 1374 | 1281 | 633 | 1671 | 12388 | 11789 | 11705 | 12146 | 11503 | 12277 |
| Foxlike silence | CB0437 | 1348 | 621 | 657 | 630 | 666 | 1225 | 1251 | 1088 | 1302 | 12073 | 12066 |
| Foxlike silence | CB0186 | 616 | 1107 | 12448 | 604 | 11829 | 1273 | 1718 | 1165 | 11254 | 12394 | 12320 |
| Foxlike silence | CB0502 | 623 | 621 | 11704 | 1635 | 636 | 11859 | 12003 | 12269 | 11783 | 11956 | 11994 |
| Foxlike silence | CB2820 | 1324 | 660 | 1311 | 631 | 12099 | 12426 | 9624 | 1256 | 12633 | 11839 | 12031 |
| Foxlike silence | CB0489 | 626 | 1312 | 627 | 639 | 12428 | 12419 | 1257 | 11832 | 9409 | 12195 | 12764 |
| Foxlike silence | CB0900 | 1357 | 622 | 633 | 629 | 675 | 636 | 11622 | 1252 | 12640 | 12414 | 12733 |
| Foxlike silence | CB0737 | 651 | 11746 | 665 | 11418 | 11852 | 640 | 1082 | 7796 | 1174 | 11877 | 12002 |
| Foxlike silence | CB0923 | 656 | 640 | 678 | 624 | 12472 | 1210 | 11840 | 12549 | 12089 | 12028 | 12007 |
| Foxlike silence | CB0479 | 1512 | 638 | 635 | 950 | 1131 | 11750 | 1252 | 11924 | 11963 | 12581 | 11527 |
| Foxlike silence | CB0587 | 12503 | 619 | 1589 | 632 | 11840 | 1351 | 12317 | 12252 | 1666 | 11810 | 11992 |
| Foxlike silence | CB0797 | 628 | 638 | 644 | 691 | 631 | 12649 | 1101 | 11466 | 12033 | 11564 | 12696 |
| Foxlike silence | CB0707 | 663 | 11697 | 623 | 627 | 666 | 632 | 12006 | 12509 | 12540 | 12633 | 11939 |
| Foxlike silence | CB0137 | 12525 | 663 | 1141 | 636 | 672 | 1214 | 633 | 11859 | 1316 | 12497 | 1343 |
| Foxlike silence | CB0014 | 618 | 640 | 656 | 626 | 1807 | 12356 | 1096 | 1376 | 12541 | 12457 | 1345 |
| Foxlike silence | CB0552 | 11913 | 628 | 636 | 11910 | 673 | 1305 | 11746 | 12375 | 11880 | 11996 | 12013 |
| Foxlike silence | CB0238 | 11379 | 646 | 1145 | 643 | 624 | 10810 | 1403 | 12543 | 1588 | 11878 | 12355 |
| Foxlike silence | CB1779 | 681 | 630 | 615 | 10793 | 647 | 1318 | 1391 | 1419 | 12431 | 12214 | 11762 |
| Foxlike silence | CB2115 | 1265 | 1395 | 629 | 634 | 1473 | 1234 | 11995 | 1223 | 12381 | 11821 | 12327 |
| Foxlike silence | CB1475 | 627 | 626 | 621 | 659 | 622 | 645 | 12192 | 9208 | 12132 | 1404 | 11441 |
| Foxlike silence | CB0917 | 624 | 620 | 631 | 1317 | 655 | 1226 | 12027 | 11705 | 1406 | 12751 | 12546 |
| Foxlike silence | CB2385 | 623 | 641 | 1107 | 619 | 1122 | 11993 | 633 | 11662 | 12535 | 12085 | 11170 |
| Foxlike silence | CB0795 | 672 | 1553 | 11775 | 1293 | 1062 | 1382 | 1151 | 10555 | 12119 | 12061 | 12589 |
| Foxlike silence | CB0468 | 628 | 637 | 633 | 626 | 1114 | 11659 | 11317 | 12307 | 12236 | 11901 | 12346 |
| Foxlike silence | CB0434 | 615 | 637 | 12041 | 1158 | 11750 | 1221 | 12090 | 10937 | 12034 | 1499 | 1505 |

| | | | | | | | | | | | | |
|---|---|---|---|---|---|---|---|---|---|---|---|---|
| Foxlike silence | CB0893 | 713 | 625 | 627 | 1241 | 11912 | 11984 | 1533 | 1289 | 1385 | 11943 | 12584 |
| Foxlike silence | CB2039 | 642 | 1360 | 626 | 642 | 1164 | 11833 | 642 | 1656 | 10847 | 12372 | 12201 |
| Foxlike silence | CB0515 | 635 | 12323 | 657 | 12526 | 656 | 1324 | 1415 | 1349 | 11950 | 8720 | 11927 |
| Foxlike silence | CB0908 | 11552 | 657 | 12554 | 633 | 640 | 1452 | 12202 | 11588 | 11900 | 1293 | 12123 |
| Foxlike silence | CB0108 | 1380 | 648 | 631 | 633 | 663 | 10293 | 11632 | 1238 | 11910 | 1663 | 11811 |
| Foxlike silence | CB2403 | 658 | 627 | 12552 | 1782 | 665 | 1423 | 1352 | 11103 | 12575 | 11554 | 12150 |
| Foxlike silence | CB0910 | 643 | 604 | 1149 | 1256 | 1233 | 11795 | 1324 | 12544 | 12183 | 11889 | 12492 |
| Foxlike silence | CB0445 | 625 | 1254 | 1636 | 11695 | 11657 | 11769 | 11925 | 12688 | 11960 | 12714 | 12479 |
| Foxlike silence | CB0604 | 645 | 12333 | 1363 | 1473 | 12218 | 931 | 1091 | 1549 | 11868 | 11024 | 11876 |
| Foxlike silence | CB3615 | 710 | 619 | 1322 | 633 | 1353 | 12225 | 12599 | 12400 | 11888 | 12318 | 12110 |
| Foxlike silence | CB0277 | 624 | 655 | 11873 | 657 | 11453 | 1362 | 10697 | 12468 | 11786 | 12674 | 1581 |
| Foxlike silence | CB0360 | 10145 | 2132 | 10876 | 661 | 630 | 11511 | 1499 | 11044 | 11859 | 12244 | 12246 |
| Foxlike silence | CB1563 | 619 | 629 | 1284 | 645 | 1290 | 12259 | 1077 | 11571 | 12512 | 11856 | 12268 |
| Foxlike silence | CB0278 | 11028 | 1288 | 1349 | 649 | 1184 | 716 | 1246 | 11919 | 12581 | 11984 | 12566 |
| Foxlike silence | CB0885 | 649 | 6804 | 1213 | 1147 | 627 | 11825 | 1179 | 657 | 1118 | 12524 | 12119 |
| Foxlike silence | CB0549 | 634 | 629 | 668 | 637 | 668 | 1344 | 1566 | 1231 | 1350 | 12479 | 11961 |
| Foxlike silence | Unknown cell type - GNG.706, GNG.656 | 644 | 628 | 640 | 643 | 11648 | 642 | 1320 | 1316 | 12386 | 12121 | 12014 |
| Foxlike silence | CB0884 | 657 | 11801 | 634 | 622 | 1276 | 11990 | 12271 | 10431 | 10511 | 11746 | 11924 |
| Foxlike silence | CB0507 | 653 | 11694 | 1430 | 637 | 1360 | 625 | 1103 | 11865 | 11708 | 12075 | 12097 |
| Foxlike silence | CB0542 | 614 | 616 | 615 | 616 | 604 | 1202 | 609 | 1085 | 12070 | 11641 | 12113 |
| Foxlike silence | CB0811 | 649 | 643 | 1355 | 624 | 1371 | 1177 | 11893 | 12069 | 1405 | 12699 | 12095 |
| Foxlike silence | CB3714 | 11957 | 636 | 609 | 1107 | 1129 | 11627 | 11749 | 12500 | 1252 | 11833 | 12454 |
| Foxlike silence | CB0879 | 661 | 644 | 661 | 627 | 647 | 1322 | 12520 | 11723 | 12092 | 12026 | 11910 |
| Foxlike silence | CB0177 | 631 | 618 | 1190 | 662 | 10946 | 1006 | 680 | 2267 | 11885 | 12445 | 11819 |
| Foxlike silence | CB0106 | 1149 | 658 | 1217 | 651 | 662 | 1160 | 669 | 11850 | 12589 | 12070 | 12538 |
| Foxlike silence | CB0581 | 612 | 1344 | 623 | 11745 | 1628 | 11717 | 12038 | 1609 | 12490 | 12012 | 12746 |
| Foxlike silence | CB0038 | 654 | 618 | 661 | 1139 | 650 | 11635 | 1266 | 11853 | 12043 | 11716 | 12412 |
| Foxlike silence | CB0493 | 1232 | 614 | 657 | 635 | 633 | 620 | 11919 | 1477 | 12016 | 12440 | 12758 |
| Foxlike silence | CB0803 | 1253 | 626 | 646 | 653 | 665 | 11116 | 1281 | 1219 | 11942 | 12063 | 12000 |
| Foxlike silence | CB0765 | 11113 | 610 | 637 | 649 | 11736 | 1116 | 1438 | 11421 | 11282 | 12045 | 11778 |
| Foxlike silence | CB3812 | 1141 | 611 | 638 | 676 | 628 | 11951 | 11564 | 1604 | 9556 | 12812 | 12154 |
| Foxlike silence | CB0775 | 11533 | 646 | 623 | 638 | 11316 | 1558 | 1368 | 11608 | 12537 | 11841 | 12613 |
| Foxlike silence | CB0239 | 12031 | 630 | 1163 | 1571 | 11604 | 11770 | 11780 | 1277 | 9637 | 12065 | 12016 |
| Foxlike silence | CB0523 | 622 | 1399 | 12283 | 1512 | 644 | 1511 | 1375 | 1574 | 12459 | 11798 | 12063 |

| | | | | | | | | | | | | |
|---|---|---|---|---|---|---|---|---|---|---|---|---|
| Foxlike silence | CB2606 | 614 | 631 | 635 | 1216 | 1141 | 1236 | 9954 | 1432 | 12044 | 11838 | 12643 |
| Foxlike silence | CB0855 | 12111 | 1240 | 666 | 11604 | 11245 | 650 | 1168 | 11741 | 1350 | 11148 | 11950 |
| Foxlike silence | CB0521 | 11554 | 8791 | 627 | 644 | 1376 | 637 | 11743 | 12129 | 11768 | 11973 | 12256 |
| Foxlike silence | CB0216 | 1440 | 630 | 644 | 626 | 1348 | 639 | 12627 | 11501 | 12496 | 1533 | 12440 |
| Foxlike silence | CB0190 | 615 | 1330 | 628 | 641 | 12310 | 12142 | 636 | 1224 | 12033 | 11951 | 11898 |
| Foxlike silence | CB0864 | 11850 | 1272 | 666 | 1223 | 626 | 636 | 1378 | 11963 | 11958 | 12250 | 12007 |
| Foxlike silence | CB0457 | 1397 | 623 | 630 | 674 | 1331 | 1338 | 1163 | 11804 | 12067 | 12665 | 12230 |
| Foxlike silence | CB0756 | 1269 | 1430 | 1211 | 625 | 1214 | 1147 | 11403 | 11932 | 11754 | 12293 | 12850 |
| Foxlike silence | CB0844 | 626 | 634 | 635 | 1235 | 635 | 639 | 12029 | 1381 | 1630 | 11681 | 12491 |
| Foxlike silence | CB1093 | 624 | 612 | 649 | 812 | 635 | 636 | 642 | 11831 | 11785 | 12030 | 1403 |
| Foxlike silence | CB0759 | 11533 | 650 | 629 | 655 | 11929 | 12191 | 11242 | 1495 | 12659 | 11572 | 11862 |
| Foxlike silence | CB0292 | 632 | 626 | 668 | 664 | 11749 | 666 | 11540 | 11750 | 1381 | 12821 | 11664 |
| Foxlike silence | CB0731 | 666 | 643 | 631 | 628 | 11689 | 1194 | 1414 | 1295 | 10876 | 12345 | 12063 |
| Foxlike silence | CB2606 | 624 | 658 | 636 | 632 | 11483 | 1579 | 11549 | 12551 | 12290 | 12296 | 11801 |
| Foxlike silence | CB1579 | 663 | 1036 | 626 | 11522 | 11866 | 1270 | 1345 | 648 | 1478 | 11670 | 11787 |
| Foxlike silence | CB0921 | 12340 | 619 | 627 | 626 | 630 | 627 | 607 | 12013 | 1500 | 11891 | 12308 |
| Foxlike silence | CB0799 | 655 | 1216 | 651 | 1085 | 11007 | 1126 | 644 | 11980 | 12031 | 1235 | 6221 |
| Foxlike silence | CB0823 | 1313 | 661 | 1421 | 1433 | 647 | 1186 | 1354 | 1308 | 1584 | 11769 | 12316 |
| Foxlike silence | CB2820 | 630 | 1214 | 11744 | 11771 | 1261 | 1041 | 1332 | 12049 | 12116 | 12463 | 12274 |
| Foxlike silence | CB2014 | 1365 | 1131 | 1226 | 1427 | 1237 | 12484 | 11819 | 12618 | 12140 | 11863 | 12184 |
| Foxlike silence | CB1470 | 1266 | 611 | 632 | 634 | 10937 | 11966 | 1231 | 11562 | 11937 | 12274 | 12161 |
| Foxlike silence | CB0731 | 628 | 668 | 1067 | 12331 | 1303 | 627 | 643 | 12043 | 1321 | 12445 | 12305 |
| Foxlike silence | CB0988 | 624 | 1111 | 634 | 697 | 633 | 11579 | 1409 | 1374 | 1272 | 12264 | 11131 |
| Foxlike silence | CB1470 | 1207 | 623 | 637 | 643 | 1311 | 1337 | 11677 | 11646 | 1126 | 11409 | 12027 |
| Foxlike silence | CB0400 | 11957 | 668 | 632 | 645 | 1478 | 11342 | 1351 | 11750 | 11914 | 1437 | 12420 |
| Foxlike silence | Unknown cell type - GNG.758 | 12362 | 638 | 666 | 1326 | 623 | 631 | 11894 | 1325 | 11797 | 11961 | 12579 |
| Foxlike silence | CB0811 | 1291 | 620 | 621 | 11645 | 1309 | 1703 | 12092 | 11728 | 10553 | 11583 | 11976 |

Table S3B. Simulated number of responding PAM-DANs to sugar GRN stimulation during silencing of select neuronal pairs or single cells.

| Experiment | Silenced neuron(s) | 100 (Hz) | 110 (Hz) | 120 (Hz) | 130 (Hz) | 140 (Hz) | 150 (Hz) | 160 (Hz) | 170 (Hz) | 180 (Hz) | 190 (Hz) | 200 (Hz) |
|---|---|---|---|---|---|---|---|---|---|---|---|---|
| Control | No silencing | 6 | 8 | 0 | 0 | 200 | 7 | 11 | 201 | 203 | 10 | 201 |
| Fox silence | CB0525 (Fox) | 0 | 0 | 0 | 0 | 0 | 0 | 0 | 0 | 0 | 0 | 0 |
| FDA silence | CB0233 | 0 | 0 | 0 | 0 | 0 | 0 | 0 | 0 | 0 | 0 | 0 |
| FDA silence | CB0546 | 0 | 202 | 0 | 0 | 200 | 7 | 0 | 7 | 199 | 196 | 205 |
| FDA silence | DNp62 | 201 | 0 | 0 | 198 | 10 | 199 | 8 | 202 | 201 | 9 | 202 |
| FDA silence | CB0272 | 0 | 0 | 0 | 201 | 200 | 7 | 0 | 7 | 199 | 196 | 205 |
| FDA silence | CB0337 | 0 | 0 | 201 | 0 | 8 | 200 | 202 | 203 | 196 | 200 | 200 |
| FDA silence | CB1514 (1) | 10 | 0 | 0 | 0 | 7 | 196 | 200 | 11 | 202 | 201 | 7 |
| FDA silence | CB1514 (2) | 6 | 0 | 7 | 0 | 7 | 8 | 0 | 11 | 200 | 200 | 202 |
| FDA silence | CB3199 | 0 | 0 | 201 | 9 | 0 | 7 | 0 | 201 | 199 | 201 | 200 |
| FDA silence | CB1025 | 197 | 9 | 0 | 6 | 0 | 6 | 8 | 203 | 199 | 194 | 205 |
| FDA silence | CB3470 | 0 | 0 | 0 | 0 | 200 | 9 | 7 | 199 | 199 | 199 | 202 |
| FDA silence | CB3573 | 9 | 0 | 0 | 10 | 200 | 3 | 8 | 200 | 9 | 200 | 202 |
| FDA silence | FDA-I | 0 | 0 | 200 | 0 | 0 | 0 | 0 | 7 | 0 | 200 | 7 |
| FDA silence | FDA-II | 0 | 0 | 0 | 0 | 0 | 0 | 0 | 0 | 0 | 0 | 0 |
| FDA silence | FDA-All | 0 | 0 | 0 | 0 | 0 | 0 | 0 | 0 | 0 | 0 | 0 |
| Foxlike silence | CB0152 | 0 | 0 | 0 | 192 | 7 | 6 | 5 | 170 | 201 | 201 | 201 |
| Foxlike silence | CB0117 | 10 | 8 | 0 | 0 | 0 | 7 | 201 | 200 | 201 | 200 | 201 |
| Foxlike silence | CB3793 | 0 | 10 | 0 | 0 | 0 | 9 | 9 | 202 | 202 | 201 | 10 |
| Foxlike silence | CB0170 | 0 | 0 | 203 | 10 | 9 | 9 | 200 | 202 | 8 | 7 | 200 |
| Foxlike silence | CB0251 | 0 | 0 | 0 | 0 | 6 | 0 | 8 | 0 | 200 | 201 | 201 |
| Foxlike silence | CB0616 | 165 | 8 | 0 | 7 | 200 | 7 | 7 | 7 | 201 | 202 | 200 |
| Foxlike silence | CB0560 | 196 | 0 | 0 | 0 | 0 | 6 | 195 | 185 | 195 | 201 | 200 |
| Foxlike silence | CB0553 | 0 | 0 | 0 | 7 | 0 | 199 | 9 | 8 | 202 | 202 | 203 |
| Foxlike silence | CB0880 | 0 | 0 | 8 | 0 | 202 | 7 | 5 | 199 | 200 | 200 | 200 |
| Foxlike silence | CB0848 | 7 | 8 | 0 | 7 | 203 | 0 | 203 | 200 | 201 | 201 | 204 |

| | | | | | | | | | | | | |
|---|---|---|---|---|---|---|---|---|---|---|---|---|
| Foxlike silence | CB0421 | 0 | 0 | 201 | 0 | 0 | 7 | 199 | 190 | 199 | 201 | 200 |
| Foxlike silence | CB0247 | 11 | 11 | 202 | 205 | 204 | 204 | 203 | 204 | 205 | 205 | 207 |
| Foxlike silence | CB0617 | 201 | 8 | 0 | 7 | 202 | 200 | 0 | 3 | 201 | 200 | 204 |
| Foxlike silence | CB0473 | 0 | 0 | 0 | 197 | 0 | 199 | 10 | 200 | 201 | 203 | 203 |
| Foxlike silence | CB0331 | 0 | 0 | 200 | 7 | 5 | 8 | 205 | 202 | 201 | 202 | 201 |
| Foxlike silence | CB0287 | 7 | 10 | 0 | 0 | 201 | 10 | 203 | 5 | 200 | 202 | 200 |
| Foxlike silence | CB0791 | 0 | 0 | 0 | 0 | 0 | 200 | 200 | 201 | 200 | 201 | 204 |
| Foxlike silence | CB0768 | 201 | 8 | 7 | 0 | 8 | 203 | 200 | 200 | 201 | 200 | 203 |
| Foxlike silence | CB0437 | 10 | 0 | 0 | 0 | 0 | 8 | 7 | 5 | 8 | 201 | 203 |
| Foxlike silence | CB0186 | 0 | 5 | 201 | 0 | 202 | 7 | 7 | 6 | 199 | 200 | 203 |
| Foxlike silence | CB0502 | 0 | 0 | 199 | 8 | 0 | 201 | 204 | 200 | 200 | 200 | 203 |
| Foxlike silence | CB2820 | 9 | 0 | 8 | 0 | 201 | 201 | 170 | 7 | 203 | 202 | 200 |
| Foxlike silence | CB0489 | 0 | 7 | 0 | 0 | 201 | 201 | 7 | 200 | 166 | 199 | 202 |
| Foxlike silence | CB0900 | 7 | 0 | 0 | 0 | 0 | 0 | 200 | 6 | 202 | 199 | 203 |
| Foxlike silence | CB0737 | 0 | 199 | 0 | 199 | 202 | 0 | 5 | 91 | 7 | 202 | 200 |
| Foxlike silence | CB0923 | 0 | 0 | 0 | 0 | 201 | 7 | 201 | 202 | 203 | 202 | 201 |
| Foxlike silence | CB0479 | 10 | 0 | 0 | 3 | 6 | 200 | 7 | 201 | 201 | 201 | 200 |
| Foxlike silence | CB0587 | 200 | 0 | 8 | 0 | 202 | 9 | 201 | 204 | 10 | 200 | 199 |
| Foxlike silence | CB0797 | 0 | 0 | 0 | 0 | 0 | 203 | 6 | 198 | 203 | 198 | 203 |
| Foxlike silence | CB0707 | 0 | 202 | 0 | 0 | 0 | 0 | 202 | 203 | 204 | 202 | 200 |
| Foxlike silence | CB0137 | 200 | 0 | 7 | 0 | 0 | 7 | 0 | 201 | 7 | 203 | 7 |
| Foxlike silence | CB0014 | 0 | 0 | 0 | 0 | 11 | 199 | 5 | 10 | 203 | 201 | 7 |
| Foxlike silence | CB0552 | 201 | 0 | 0 | 202 | 0 | 10 | 202 | 199 | 201 | 203 | 203 |
| Foxlike silence | CB0238 | 199 | 0 | 6 | 0 | 0 | 195 | 9 | 203 | 10 | 201 | 202 |
| Foxlike silence | CB1779 | 0 | 0 | 0 | 192 | 0 | 8 | 8 | 9 | 201 | 201 | 202 |
| Foxlike silence | CB2115 | 7 | 10 | 0 | 0 | 7 | 7 | 203 | 7 | 201 | 200 | 201 |
| Foxlike silence | CB1475 | 0 | 0 | 0 | 0 | 0 | 0 | 201 | 144 | 201 | 11 | 196 |
| Foxlike silence | CB0917 | 0 | 0 | 0 | 7 | 0 | 7 | 200 | 200 | 9 | 202 | 202 |
| Foxlike silence | CB2385 | 0 | 0 | 6 | 0 | 7 | 202 | 0 | 198 | 203 | 202 | 197 |
| Foxlike silence | CB0795 | 0 | 7 | 201 | 8 | 6 | 10 | 6 | 185 | 202 | 201 | 202 |
| Foxlike silence | CB0468 | 0 | 0 | 0 | 0 | 6 | 200 | 200 | 200 | 200 | 201 | 201 |
| Foxlike silence | CB0434 | 0 | 0 | 199 | 7 | 201 | 7 | 201 | 193 | 202 | 8 | 8 |
| Foxlike silence | CB0893 | 0 | 0 | 0 | 7 | 203 | 201 | 10 | 9 | 12 | 202 | 202 |

| | | | | | | | | | | | | |
|---|---|---|---|---|---|---|---|---|---|---|---|---|
| Foxlike silence | CB2039 | 0 | 7 | 0 | 0 | 7 | 203 | 0 | 9 | 193 | 200 | 202 |
| Foxlike silence | CB0515 | 0 | 202 | 0 | 201 | 0 | 8 | 8 | 8 | 201 | 122 | 200 |
| Foxlike silence | CB0908 | 200 | 0 | 204 | 0 | 0 | 8 | 200 | 201 | 201 | 7 | 203 |
| Foxlike silence | CB0108 | 11 | 0 | 0 | 0 | 0 | 184 | 201 | 7 | 202 | 9 | 200 |
| Foxlike silence | CB2403 | 0 | 0 | 200 | 8 | 0 | 10 | 12 | 196 | 202 | 199 | 203 |
| Foxlike silence | CB0910 | 0 | 0 | 6 | 7 | 7 | 199 | 10 | 202 | 202 | 202 | 203 |
| Foxlike silence | CB0445 | 0 | 8 | 10 | 199 | 200 | 200 | 201 | 203 | 201 | 203 | 202 |
| Foxlike silence | CB0604 | 0 | 200 | 8 | 8 | 201 | 3 | 5 | 7 | 200 | 199 | 200 |
| Foxlike silence | CB3615 | 0 | 0 | 9 | 0 | 7 | 200 | 202 | 202 | 200 | 203 | 202 |
| Foxlike silence | CB0277 | 0 | 0 | 199 | 0 | 200 | 10 | 188 | 200 | 200 | 201 | 7 |
| Foxlike silence | CB0360 | 182 | 9 | 190 | 0 | 0 | 199 | 9 | 195 | 201 | 201 | 201 |
| Foxlike silence | CB1563 | 0 | 0 | 9 | 0 | 7 | 201 | 6 | 199 | 202 | 200 | 202 |
| Foxlike silence | CB0278 | 196 | 7 | 7 | 0 | 7 | 0 | 7 | 198 | 203 | 202 | 202 |
| Foxlike silence | CB0885 | 0 | 63 | 7 | 6 | 0 | 199 | 7 | 0 | 6 | 203 | 202 |
| Foxlike silence | CB0549 | 0 | 0 | 0 | 0 | 0 | 7 | 10 | 6 | 9 | 202 | 203 |
| Foxlike silence | Unknown cell type - GNG.706, GNG.656 | 0 | 0 | 0 | 0 | 200 | 0 | 7 | 11 | 201 | 202 | 201 |
| Foxlike silence | CB0884 | 0 | 199 | 0 | 0 | 8 | 200 | 199 | 188 | 187 | 200 | 201 |
| Foxlike silence | CB0507 | 0 | 201 | 6 | 0 | 9 | 0 | 7 | 202 | 200 | 200 | 200 |
| Foxlike silence | CB0542 | 0 | 0 | 0 | 0 | 0 | 7 | 0 | 6 | 203 | 200 | 202 |
| Foxlike silence | CB0811 | 0 | 0 | 10 | 0 | 9 | 7 | 201 | 201 | 8 | 202 | 201 |
| Foxlike silence | CB3714 | 200 | 0 | 0 | 5 | 6 | 198 | 201 | 202 | 7 | 200 | 203 |
| Foxlike silence | CB0879 | 0 | 0 | 0 | 0 | 0 | 7 | 202 | 200 | 203 | 201 | 201 |
| Foxlike silence | CB0177 | 0 | 0 | 6 | 0 | 196 | 5 | 0 | 9 | 200 | 202 | 201 |
| Foxlike silence | CB0106 | 6 | 0 | 6 | 0 | 0 | 7 | 0 | 201 | 203 | 200 | 202 |
| Foxlike silence | CB0581 | 0 | 7 | 0 | 202 | 10 | 200 | 201 | 10 | 203 | 200 | 200 |
| Foxlike silence | CB0038 | 0 | 0 | 0 | 6 | 0 | 200 | 7 | 199 | 202 | 200 | 203 |
| Foxlike silence | CB0493 | 5 | 0 | 0 | 0 | 0 | 0 | 202 | 8 | 201 | 202 | 203 |
| Foxlike silence | CB0803 | 7 | 0 | 0 | 0 | 0 | 195 | 7 | 7 | 202 | 203 | 201 |
| Foxlike silence | CB0765 | 194 | 0 | 0 | 0 | 201 | 6 | 8 | 198 | 196 | 200 | 199 |
| Foxlike silence | CB3812 | 4 | 0 | 0 | 0 | 0 | 201 | 198 | 8 | 160 | 204 | 203 |
| Foxlike silence | CB0775 | 199 | 0 | 0 | 0 | 199 | 10 | 8 | 199 | 201 | 199 | 202 |
| Foxlike silence | CB0239 | 200 | 0 | 6 | 10 | 200 | 201 | 200 | 7 | 170 | 200 | 202 |
| Foxlike silence | CB0523 | 0 | 7 | 201 | 10 | 0 | 8 | 9 | 7 | 202 | 199 | 202 |
| Foxlike silence | CB2606 | 0 | 0 | 0 | 7 | 6 | 7 | 177 | 9 | 202 | 200 | 200 |

| | | | | | | | | | | | | |
|---|---|---|---|---|---|---|---|---|---|---|---|---|
| Foxlike silence | CB0855 | 202 | 7 | 0 | 200 | 198 | 0 | 6 | 200 | 10 | 195 | 200 |
| Foxlike silence | CB0521 | 202 | 124 | 0 | 0 | 9 | 0 | 201 | 201 | 201 | 203 | 202 |
| Foxlike silence | CB0216 | 8 | 0 | 0 | 0 | 7 | 0 | 204 | 201 | 201 | 9 | 204 |
| Foxlike silence | CB0190 | 0 | 7 | 0 | 0 | 202 | 200 | 0 | 7 | 200 | 202 | 201 |
| Foxlike silence | CB0864 | 199 | 8 | 0 | 7 | 0 | 0 | 7 | 201 | 202 | 203 | 200 |
| Foxlike silence | CB0457 | 7 | 0 | 0 | 0 | 11 | 7 | 5 | 201 | 200 | 202 | 201 |
| Foxlike silence | CB0756 | 7 | 10 | 6 | 0 | 7 | 7 | 197 | 201 | 202 | 202 | 202 |
| Foxlike silence | CB0844 | 0 | 0 | 0 | 8 | 0 | 0 | 200 | 10 | 12 | 198 | 202 |
| Foxlike silence | CB1093 | 0 | 0 | 0 | 0 | 0 | 0 | 0 | 201 | 200 | 202 | 9 |
| Foxlike silence | CB0759 | 198 | 0 | 0 | 0 | 202 | 203 | 199 | 10 | 202 | 200 | 200 |
| Foxlike silence | CB0292 | 0 | 0 | 0 | 0 | 201 | 0 | 199 | 200 | 8 | 203 | 199 |
| Foxlike silence | CB0731 | 0 | 0 | 0 | 0 | 200 | 7 | 11 | 8 | 196 | 203 | 201 |
| Foxlike silence | CB2606 | 0 | 0 | 0 | 0 | 201 | 8 | 201 | 201 | 202 | 203 | 201 |
| Foxlike silence | CB1579 | 0 | 6 | 0 | 199 | 201 | 8 | 10 | 0 | 7 | 200 | 200 |
| Foxlike silence | CB0921 | 201 | 0 | 0 | 0 | 0 | 0 | 0 | 203 | 9 | 201 | 203 |
| Foxlike silence | CB0799 | 0 | 9 | 0 | 5 | 194 | 6 | 0 | 201 | 201 | 7 | 57 |
| Foxlike silence | CB0823 | 10 | 0 | 10 | 9 | 0 | 6 | 9 | 9 | 10 | 201 | 205 |
| Foxlike silence | CB2820 | 0 | 7 | 199 | 201 | 7 | 6 | 8 | 201 | 200 | 202 | 202 |
| Foxlike silence | CB2014 | 8 | 6 | 9 | 7 | 9 | 202 | 200 | 202 | 202 | 202 | 201 |
| Foxlike silence | CB1470 | 9 | 0 | 0 | 0 | 194 | 200 | 7 | 200 | 201 | 205 | 201 |
| Foxlike silence | CB0731 | 0 | 0 | 6 | 201 | 10 | 0 | 0 | 201 | 10 | 201 | 203 |
| Foxlike silence | CB0988 | 0 | 6 | 0 | 0 | 0 | 196 | 8 | 8 | 7 | 202 | 194 |
| Foxlike silence | CB1470 | 9 | 0 | 0 | 0 | 8 | 7 | 199 | 199 | 6 | 198 | 202 |
| Foxlike silence | CB0400 | 199 | 0 | 0 | 0 | 8 | 199 | 7 | 201 | 199 | 8 | 201 |
| Foxlike silence | Unknown cell type - GNG.758 | 201 | 0 | 0 | 8 | 0 | 0 | 201 | 8 | 202 | 200 | 201 |
| Foxlike silence | CB0811 | 8 | 0 | 0 | 198 | 7 | 9 | 202 | 200 | 191 | 200 | 201 |
